# Pivotal role of Disrupted-in-Schizophrenia 1 (DISC1) in cardiac resilience to ischemic stress

**DOI:** 10.1101/2022.06.09.494639

**Authors:** Gurdeep Marwarha, Maria J. Pinho, Nathan R Scrimgeour, Katrine Hordnes Slagsvold, Alexander Wahba, Ragnhild E Røsbjørgen, Francisco J. Enguita, Kun Yang, Koko Ishizuka, Carlos Andrés Acosta Casas, Rajeevkumar Raveendran Nair, Rita Brekke, Kristine Pettersen, Geir Bjørkøy, Akira Sawa, Morten A Høydal

## Abstract

Molecular targets that contribute to post-myocardial infarction heart failure remains elusive. Here we studied the global transcriptional landscape of human inducible pluripotent stem cell-derived cardiomyocytes under simulated ischemia at the whole genome level in an unbiased manner. Unexpectedly, we identified Disrupted in Schizophrenia 1 (DISC1), which has been almost exlusively studied in neurodevelopment/neurosignaling, as a key molecule. Reduced DISC1 levels increase infarct size in mouse hearts, increases cTnT release following surgery in coronary artery bypass grafting patients and decrease the survival of human AC16 cardiomyocytes exposed to ischemic conditions by disrupting cardioprotective pathways. Mechanistically, the loss of DISC1 interaction with Glycogen Synthase Kinase 3 Beta (GSK3β) heightens GSK3β activity, promoting cell death. Conversely, increasing DISC1 levels enhances cardioprotective signaling by maintaining the DISC1-GSK3β interaction. Although DISC1 is known to interact with many proteins as an intracellular hub, we have identified that its specific interaction with GSK3β is crucial for cardioprotection. These data also indicate that a key molecule involved in brain health and disease also contributes to cardiac health and disease, supporting the idea that neuropsychiatric conditions are systemic.

**Summary:** Through an unbiased screening at the whole genome level, we underscore that DISC1 is a key molecule for cardiomyocyte resilience post-myocardial infarction. DISC1 has been studied almost exclusively in neuroscience, involving in multiple cellular mechanisms. In contrast, we highlight DISC1’s interaction with GSK3β as specific mechanism that drives cardioprotection. We also report DISC1’s implication in cardiac health following coronary artery bypass grafting.

## Introduction

Ischemic heart disease, the leading cause of death worldwide,^1^ occurs when reduced blood flow causes oxygen deprivation in the heart muscle. This oxygen deprivation, or ischemia, can lead to myocardial infarction, commonly known as a heart attack.^2^ Following a myocardial infarction, the activation of various intracellular signaling pathways triggers pathological heart remodeling, further driving disease progression.^3^ Numerous efforts have been made to understand these pathological processes, in the hope that better understanding may protect the heart against ischemia, but there is still a major knowledge gap.^2,4–6^

To address this challenge, we initially looked at the global transcriptional landscape through next-generation RNA sequencing in human inducible pluripotent stem cell-derived cardiomyocytes (hiPS-cardiomyocytes) subjected to hypoxic stress. Exploring this landscape allowed an identification of novel key transcripts involved in the cardiomyocyte response to hypoxia. Unexpectedly, through an unbiased bioinformatics analysis, disrupted in schizophrenia 1 (DISC1) emerged as the transcript in cardiomyocytes subjected to hypoxic conditions that exhibited the most substantial changes. This robust alteration underscores the potential significance of DISC1 in the cellular response to hypoxia. DISC1 has been studied as an intracellular scaffold protein that plays a key role in neurodevelopment^7,8^ In a large Scottish pedigree, the disruption of this molecule is likely to underlie aggregation of psychotic and mood disorders among the family members^9–11^. Accordingly, DISC1 has been studied in the context of neurodevelopment, neurosignaling, and their implications in neurodevelopmental psychiatric disorders^7,12^. To the best of our knowledge, DISC1 has not been studied in cardiology thus far.

Compared with the general population, patients with psychotic disorders such as schizophrenia are prone to premature death, with a one-to-two-decade reduction in life expectancy^13,14^. Ischemic heart disease is a major reason for this premature mortality^15–17^, although antipsychotic medications can also be a contributing factor. Some recent studies have reported deteriorated cardiac function measured by echocardiography and MRI in patients with schizophrenia^18,19^. Myocardial T1 mapping (proton spin-lattice relaxation time), which enables detection of myocardial edema, infarction, ischemia, cardiomyopathies and diffuse fibrosis^20^, further suggests that an early diffuse fibroinflammatory myocardial process in patients with schizophrenia is independent of established cardiovascular disease risk factors and could contribute to excess cardiovascular mortality.^19^

Motivated by the initial global transcriptional mapping that highlighted DISC1, we hypothesized that DISC1 may play a significant role beyond its well-established effects in the brain, specifically addressing its causal mechanism in heart hypoxia. In the present study, we demonstrate that DISC1 plays a crucial regulatory role in heart hypoxia, protecting against cardiac damage caused by ischemic events, such as myocardial infarction. This protective effect is mediated through the regulation of GSK3β activity and its downstream cascade, which ultimately reduces apoptotic cell death.

## Results

### Global transcriptional profiling and unbiased bioinformatics highlight DISC1 in cardiac hypoxia

In the initial phase of this project, we employed an unbiased approach to explore unknown signaling cascades in heart hypoxia. We examined the global transcriptional landscape through RNA sequencing (RNA-seq) of human inducible pluripotent stem cell-derived cardiomyocytes (hiPS-cardiomyocytes) that were cultured for 48 hours in a starvation medium and exposed to either 1% O_2_ or normoxia as a control. This analysis revealed 1232 significantly regulated coding transcripts (adjusted P < 0.05) between hypoxia and normoxia (***Supplements Appendix 1***). To further narrow down the potential novel candidates, we applied stringent statistical criteria with a cut-off -Log_10_ P value < 10E^-20^ and -Log_2_ fold change (FC) cutoff <-2 or > +2. This approach identified a subset of transcripts that met these criteria: Coiled-Coil Domain Containing 134 (CCDC134), Thyroid Hormone Receptor Alpha (THRA), CKLF Like MARVEL Transmembrane Domain Containing 5 (CMTM5), DISC1, Synaptotagmin 2 (SYT2), Isoprenylcysteine Carboxyl Methyltransferase (ICMT), Zinc Finger Protein 142 (ZNF142), and Thymopoietin (TMPO) (**Fig 1 A-B**).

**Fig. 1.**
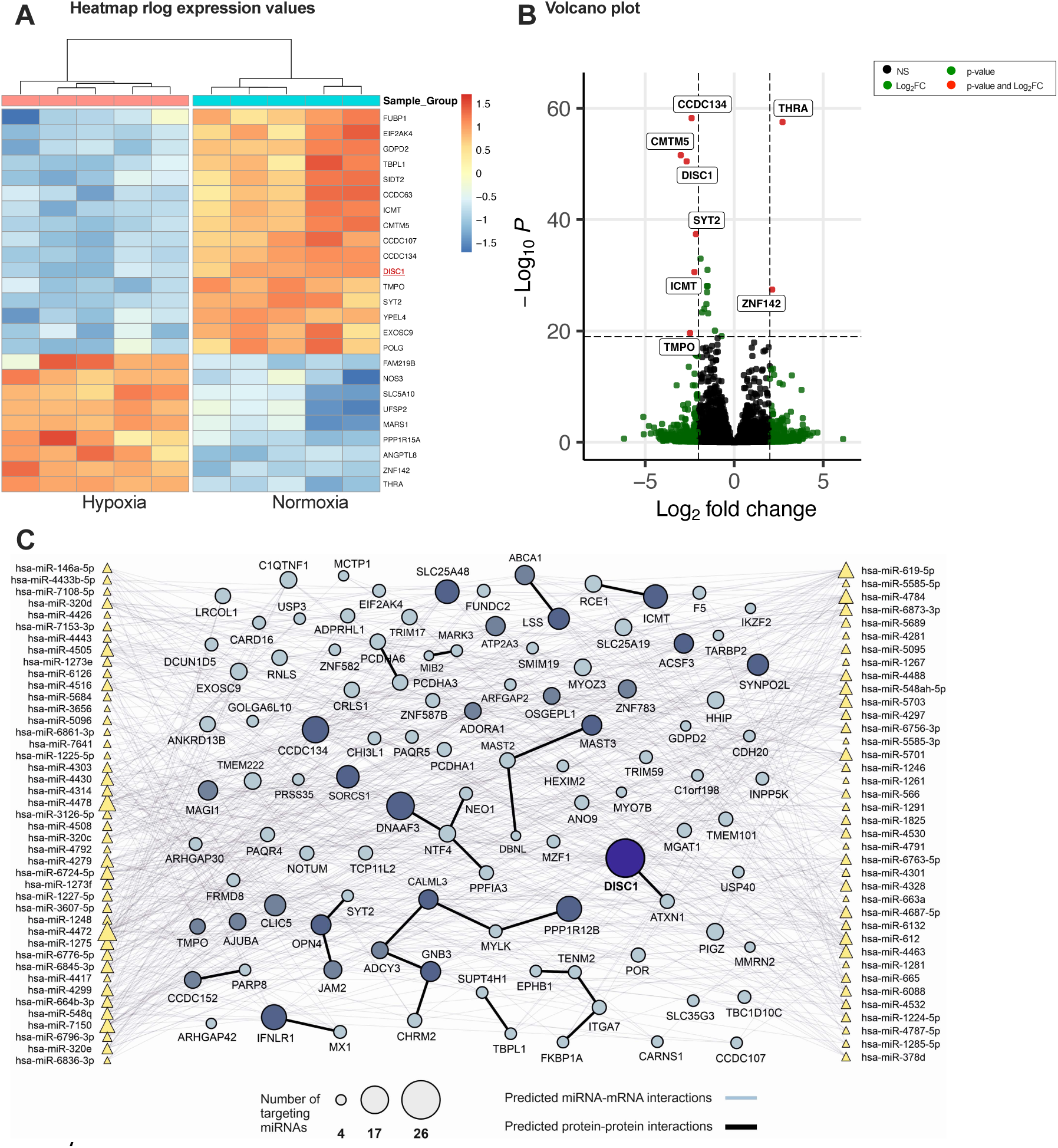
Discovering DISC1 by transcriptomics. **A,** Heatmap of RNA sequencing data of the coding RNA with a cutoff value of 10E^-10^ and a Log_2_FC cutoff >2. **B**, Volcano plot displaying all regulated coding mRNA transcripts including statistical analyses with cut-off -Log_10_ P value < 10E^-20^ and -Log_2_ fold change value ± 2, mitochondrial mRNA are left out for simplicity in A-B (all data included in online supplements). **C**, Regulatory networks in hypoxia-stimulated hIPSC-cardiomyocytes relationships between upregulated microRNAs (miRs) and their downregulated mRNA targets. Upregulated miRNAs are depicted as triangles, whose sizes are related to the number of targets; mRNAs are represented by circles; black lines represent the interaction between proteins encoded by each gene extracted from STRING (combined score >0.8) (Supporting data in Excel file, E3).

To further explore the global transcriptional response to hypoxia in cardiomyocytes, we expanded our bioinformatics analyses by generating regulatory networks using input from the RNA-seq data on both coding and non-coding microRNA transcripts. These analyses focused on negative correlations between overexpressed microRNAs (miRs) and their putative downregulated mRNA targets. Our analyses identified a total of 774 miR-mRNA pairs. DISC1 emerged as the top candidate gene, standing out with the highest number of miR-mRNA interactions, being targeted by 26 specific miRs. Among the other transcripts determined from the statistical cut-off, we found 17 miR-mRNA interactions for CCDC134, 14 for ICMT, 8 for TMPO, 4 for SYT2, and none for ZNF142, THRA, and CMTM5. DISC1’s prominence underscores its importance among all 1232 significantly regulated coding transcripts (**Fig. 1C *and extended data in supplements Appendix 2 and 3)*.**

In the RNA sequencing data, DISC1 displayed a downregulation with a Log2FC of -2.67 under hypoxic conditions (P-value = 3.41E-51) (**Fig. 1A-B** and ***supplements Fig. S1, Extended Data in Excel file, Appendix 1***). The robust significant change of DISC1 transcript following hypoxia combined with the number of miRs targeting DISC1, suggests a complex regulatory mechanism, highlighting its potential role in the adaptive response to hypoxic stress. This initial unbiased discovery of DISC1 sparked our interest in understanding its role, which has not been previously described in the context of the heart.

### Enrichment of cardiovascular risk gene products among DISC1 interactors

To understand if DISC1 could be an interesting candidate for further analyses we explore the potential involvement of DISC1 in cardiovascular-related conditions from a genetic perspective. From earlier studies in neuroscience, DISC1 is described to acts as a scaffold protein, interacting with numerous proteins to form complex intracellular signaling networks.^7,8^ We hypothesized that risk gene products (proteins) associated with cardiovascular conditions may be enriched among DISC1 interactors. To test this, we first identified risk genes for traits linked to cardiologic diseases, malformations, or pathologies (***Supplements Table S1, Data S1***) using data from the NHGRI-EBI GWAS Catalog (criteria: P-value < 1E-8 and odds ratio > 1). A chi-square test was then performed to determine whether these risk gene products were overrepresented among DISC1 interactors, compared to non-interacting genes in the human genome.^21^ Our analysis revealed a significant enrichment of cardiovascular risk gene products among DISC1 interactors (P-value = 2.8E-17). These findings suggest that downregulation of DISC1 may play a critical role in hypoxia, increasing the heart’s vulnerability to ischemic damage.

Taken together, with a strong support from these unbiased bioinformatic data, we decided to conduct cell biological experiments to validate this notion and address the mechanism of how DISC1 may play this role in the heart and cardiomyocytes.

### Biphasic response in DISC1 expression: time and severity-dependent alterations in cardiac ischemia and cardiomyocyte hypoxia

To confirm the presence of DISC1 in cardiomyocytes, as well as the regulation of DISC1 in different models and clinically relevant conditions of ischemic heart disease, we first measured mRNA expression in a rat model of myocardial infarction. Compared to sham-operated rat hearts, DISC1 was significantly upregulated in viable myocardium remote from the infarct area, whereas in the ischemic border zone region, we found unaltered expression of DISC1 transcript (**Fig. 2A**). Analyses in human cardiac left ventricle biopsies from post-myocardial infarction coronary artery bypass grafting (CABG) patients displayed a tendency towards increased DISC1 expression in the remote viable myocardium (**Fig. 2B**), indicating increased DISC1 in response to cardiac ischemia.

**Fig. 2.**
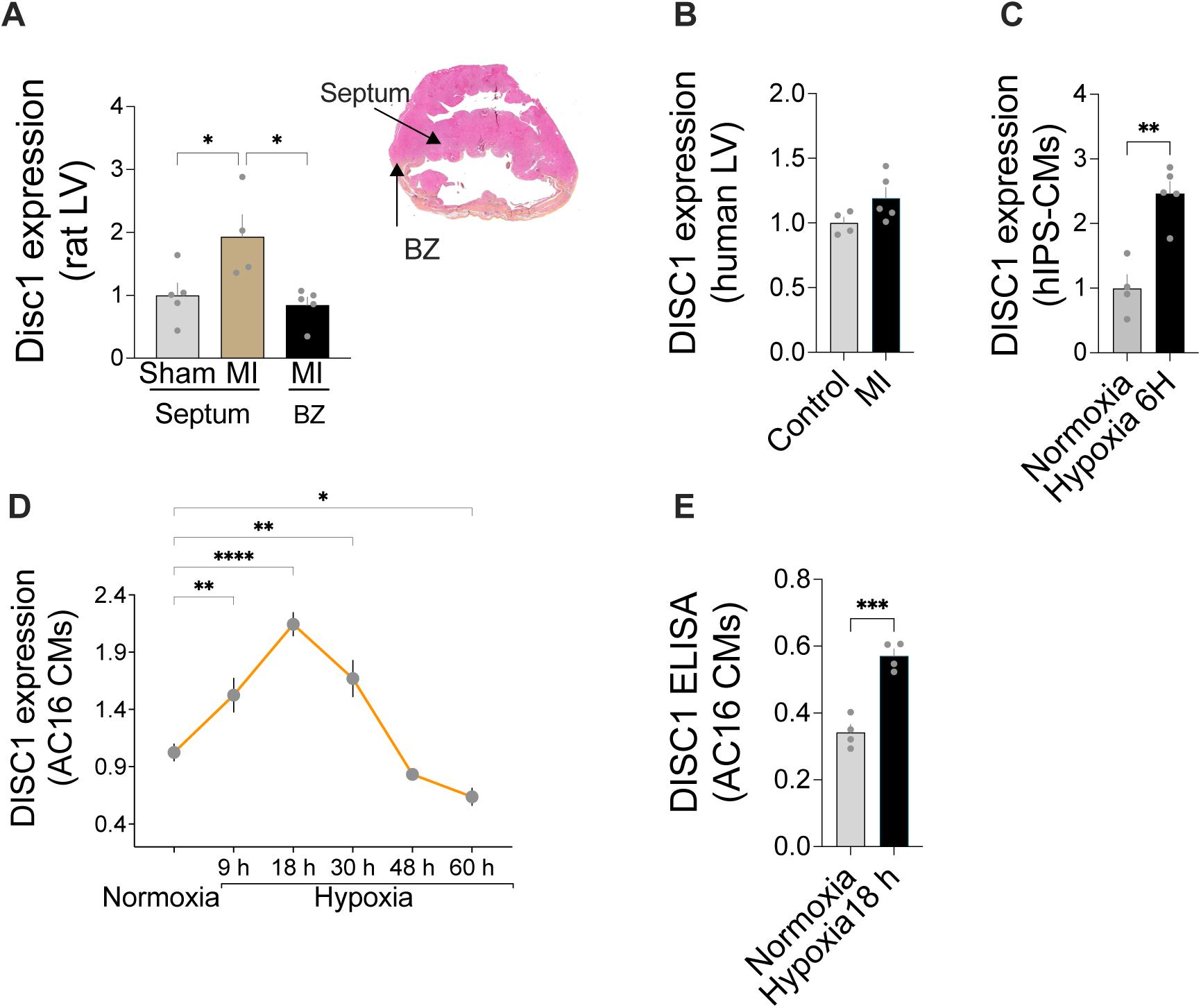
Expressional regulation of DISC1 in myocardial infarction and cardiomyocyte hypoxia. **A**, DISC1 mRNA expression changes in post-myocardial infarction (MI) rat hearts from sham-operated animals and regional areas distant from the MI (sham, N=5; MI septum, N=4: MI border zone (BZ), N=5). Zones of harvesting displayed by exemplary rat heart with MI. **B**, DISC1 mRNA expression in CABG patient biopsies from the non-infarcted area of the left ventricle of post-MI heart failure patients (N=5) vs. non-MI controls (N=4). **C**, Quantification of DISC1 mRNA transcripts in hiPS-CM following 6 hours of hypoxia (1% O_2_). **D**, Timeline of DISC1 mRNA expression in human AC16 CMs following normoxia and hypoxia. **E**, DISC1 total protein expression following 18 hours of hypoxia in AC16 CMs. Data presented as fold change(A-D); E is expressed as absorbances measured at λ_450_ (450 nm), data presented as mean ± s.e.m. P values are indicated by : *\* P<0.05, ** P<0.01, *** P<0.001, **** P<0.0001*. Statistical analyses in A performed by one-way ANOVA and Tukey’s post hoc test; B, C and E performed by unpaired t-test; D, performed by one-way ANOVA compared to normoxia. All analyses performed using two-tailed analyses.

Previous studies from neural cells have reported that the expression of DISC1 is significantly downregulated following hypoxia.^22^ These data are in line with our initial discovery of the regulation of DISC1 in cardiomyocytes, where we found that DISC1 was significantly reduced in hiPS-cardiomyocytes stimulated by long-term exposure to hypoxia (48 hours, 1% O_2_) (**Fig 1A-B**). Given this contradiction in DISC1 expression compared to less ischemic impacted areas of the heart, we tested shorter timepoints of hypoxia-stimulated cardiomyocytes in culture. Indeed, 6 hours of hypoxia in hiPS-cardiomyocytes significantly increased DISC1 (**Fig. 2C**). The hypoxic response of DISC1 expression was further tested in human AC16 cardiomyocytes, which confirmed a time-dependent response, with DISC1 expression increasing up to 18 hours and thereafter declining to levels below the normoxia baseline at 48 hours and 60 hours. (**Fig. 2D-E**). Our data suggest that the initial upregulation of DISC1 expression within the first hours of ischemic/hypoxic stress in cardiomyocytes acts as a compensatory mechanism, potentially conferring protection to the cardiomyocytes and enhancing their resistance to hypoxic conditions.

Conversely, the maladaptive response of cardiomyocytes, characterized by reduced or diminished DISC1 expression during prolonged ischemic and/or hypoxic stress, may exacerbates ischemic damage to the heart and cardiomyocytes. Therefore, to further investigate the role of DISC1 in ischemic damage, we examined whether downregulation of DISC1 influences myocardial infarction size.

### Downregulation of DISC1 increases myocardial infarction size

To test whether downregulation of DISC1 causes greater ischemic damage, we used a mouse model with a DISC1-locus deletion (DISC1-locus impairment (DISC1-LI), as previously described.^8^ We found that downregulation of DISC1 caused significantly larger infarct size following ischemia-reperfusion in a Langendorff coronary perfusion infarct model of thirty minutes of global ischemia followed by 2 hours of reperfusion in isolated DISC1-Li mouse hearts vs. wild type (WT) mouse hearts. DISC1-Li mice displayed larger infarct sizes in both male (130% larger) and female mice (73% larger) (**Fig. 3A-C**). These data strengthen the hypothesis that downregulation of DISC1 indeed exacerbates ischemic damage.

**Fig. 3.**
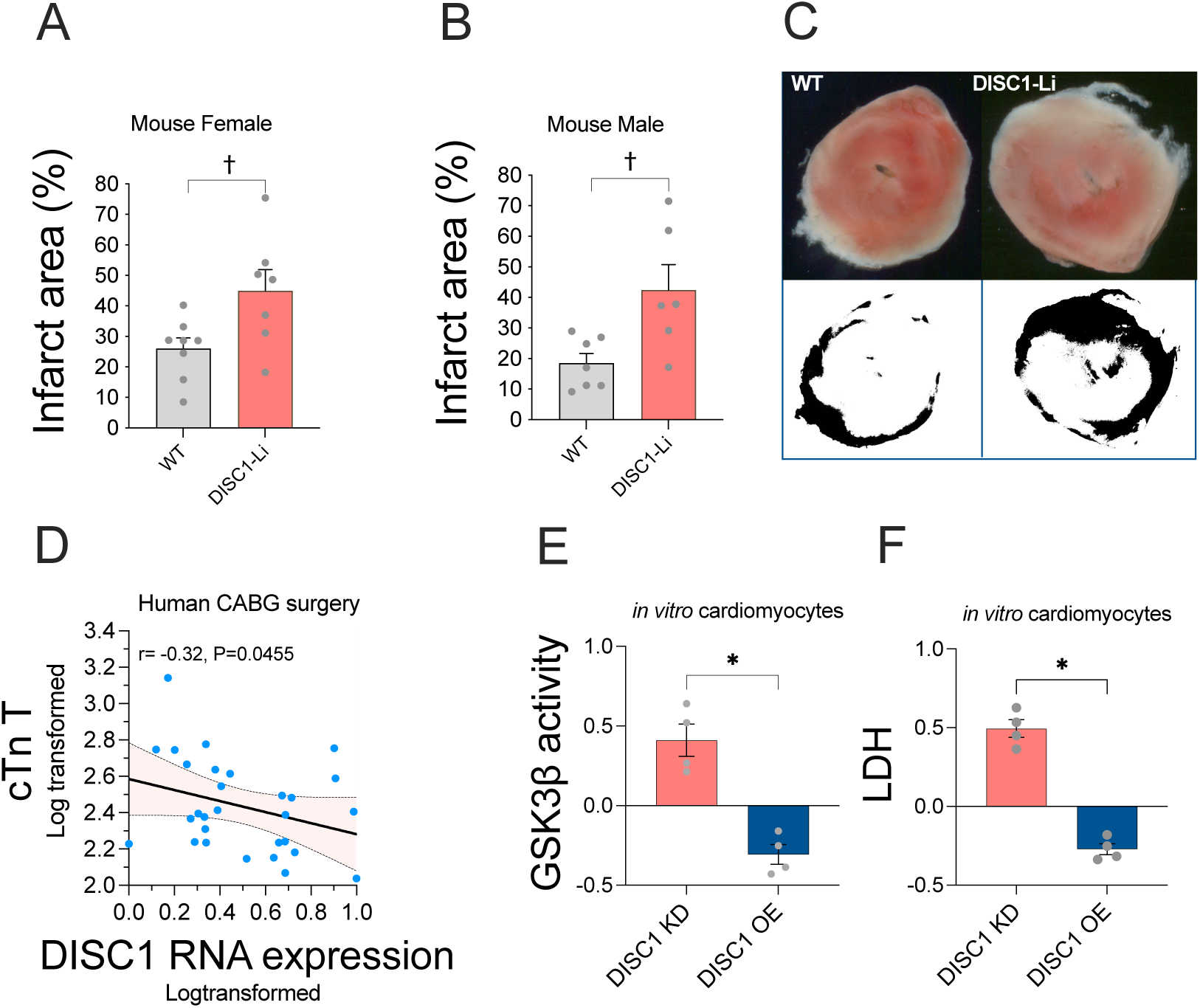
Reduced DISC1 expression increases cardiac damage following ischemic and hypoxic stress, in mouse models, Coronary Artery Bypass Grafting (CABG) patients and in vitro cardiomyocyte models. **A-B**, Myocardial infarct size in WT and DISC1-Li mice after 30 minutes of global ischemia followed by 1 h 40 m of reperfusion in male mice (WT, n=7/DISC1-Li, n=6) and female mice (WT, n=7/DISC1-Li, n=7). **C**, Exemplary images from WT and DISC-Li Male. The black/white image shows the infarct area in black. **D**, Data from CABG patients showing the delta release of cardiac Troponin T (cTnT) from preoperative to 24 hours postoperative, compared to DISC1 RNA expression in blood measured preoperatively (N=29). Figures **E-F** represent experiments in in AC16 cardiomyocytes to determine the effect of both DISC1 silencing using short hairpin (sh)RNA pLKO.1-DISC1 (**DISC1 KD**) compared to respective pLKO.1-empty vector or by overexpression of DISC1 using pRK5-DISC1 overexpression vector (DISC1 OE) compared to respective pRK5-EV. All data are presented in relation to the fractional difference from respective Mock (zero line). **E**, GSK3β activity and **F**, Lactate dehydrogenase released from cardiomyocytes to cytosol. Statistical analyzes in A-B are performed using two-tailed t-test († *P<0.05,* difference between WT and DISC1-Li mice). Data in panel D exhibited a lognormal distribution and were log-transformed for both cardiac Troponin T (cTnT) and DISC1 RNA expression. Correlation was analysed using one-tailed Pearson correlation. Statistical analyses in E-F were performed using using two-tailed t-test to assess the difference in response compared to respective Mock between DISC1 KD versus DISC1 OE. Statistical difference displayed by: *\* P<0.05, ** P<0.01, *** P<0.001*.

### Low expression levels of DISC1 RNA in blood correletes with increased release of cardiac troponin T (cTnT) following CABG surgery

Based on the significantly increased infarct size in mice with downregulated DISC1 we aimed to verify our findings in a clinical study from CABG patients, a setting where all patients underwent peroperative cardiac ischemia-reperfusion due to aortic cross-clamping, as reflected by postoperative release of cTnT.^23–25^ Therefore, we investigated whether there was a correlation between low DISC1 RNA expression in blood and increased cTnT release from pre-to 24 hours post-CABG surgery, as a marker of ischemia-reperfusion injury. Our analysis revealed that patients with lowest DISC1 RNA levels displayed highest cTnT release 24 hours post-surgery (Pearson correlation, r = -0.32, P = 0.045, **Fig. 3D**). These findings validate our hypothesis and emphasize the clinical significance of DISC1 in enhancing cardiac resilience to ischemia.

### Downregulation of DISC1 exacerbates cardiomyocyte vulnerability to hypoxic stress trough activation of GSK3β while overexpression of DISC1 reverses these effects

Oxygen and nutrient deprivation triggers metabolic acidosis, calcium overload, and reactive oxygen species (ROS), leading to mitochondrial depolarization, swelling, and opening of the mitochondrial permeability transition pore (mPTP),^26,27^ ultimately resulting in cell death.^28–30^ To gain deeper mechanistic insights into the impact of modified DISC1 expression on cardiomyocyte death, we employed shRNA-DISC1 for silencing and a DISC1 expression vector for overexpression. We found that silencing of DISC1 significantly sensitized cardiomyocytes to 18 hours of hypoxic stress and increased the activity of GSK3β (**Fig. 3E**), which culiminated in increased cell death assayed by lactate dehydrogenase (LDH) release (**Fig. 3F**). Further analyses verified activation of established downstream signalling processes, including BAX’s interaction with the outer mitochondrial membrane (OMM), resulting in membrane permeabilization, CytC release, and the activation of caspase-3 (***Supplements Fig. S2-5***). Importantly, overexpression of DISC1 inhibited the activation of GSK3β (**Fig. 3E**) and reversed downstream cell death signaling processes (**Fig. 3F**, ***Supplements Fig. S2-5***). These data confirm an important role of DISC1 in suppressing GSK3β activity, leading to enhanced cardiomyocyte resistance against hypoxia-induced cell death^31^.

### Causative link between DISC1-GSK3β interaction for the regulation of cell death signaling?

To this point we have comprehensively investigated the role of DISC1; we meticulously examined its impact on GSK3β activation, and confirmed known signalling processes of hypoxia induced cell death which were amplified with silencing of DISC1 and mitigated by overexpression of DISC1. We also confirmed that there exist a direct interaction between DISC1 and GSK3β by Co-IP in human AC16 cardiomyocytes (***Supplements Fig S6***). However, a lingering question remained: can the enhanced GSK3β activity observed in cells with downregulated DISC1, and conversely, the reduced GSK3β activity in cells with overexpressed DISC1, be attributed to a direct interaction between DISC1 and GSK3β? To address this question, we designed additional experiments where we assessed the effect of interrupting the binding between DISC1 and GSK3β.

First, we designed a new overexpression vector with a triple mutant and replaced Phenylalanines (F) with Alanine (A) at the sites (F210, F212 and F206) in the full length DISC1 (3AAA mutant), that would not allow for DISC1-GSK3β interaction, based on a previous report that the residues 193−236 from the full-length protein are the most potent GSK3β binding region of DISC1.^32^ To avoid an effect of the DISC1 shRNA on the overexpression product, silent nucleotide mutations at the target sequence of the DISC1 shRNA were included into the overexpression vector, thus hindering the silencing of the expressed 3AAA mutant or wild type DISC1. Using this approach the expressed product of DISC1 should be a DISC1-3AAA mutant resistant to RNAi (DISC1-ΔGSK3β^R^) or full length (FL) wild type DISC1 resistant to RNAi (DISC1-FL^R^). Then, we verified the DISC1-ΔGSK3β^R^ effect for DISC1-GSK3β interaction using Co-Immunoprecipitation (Co-IP). We found that the interaction between for DISC1-GSK3β was significantly reduced in the DISC1-ΔGSK3β^R^, confirming that mutating the F210, F212 and F206 sites hinder interaction (**Fig. 4A**).

**Fig. 4.**
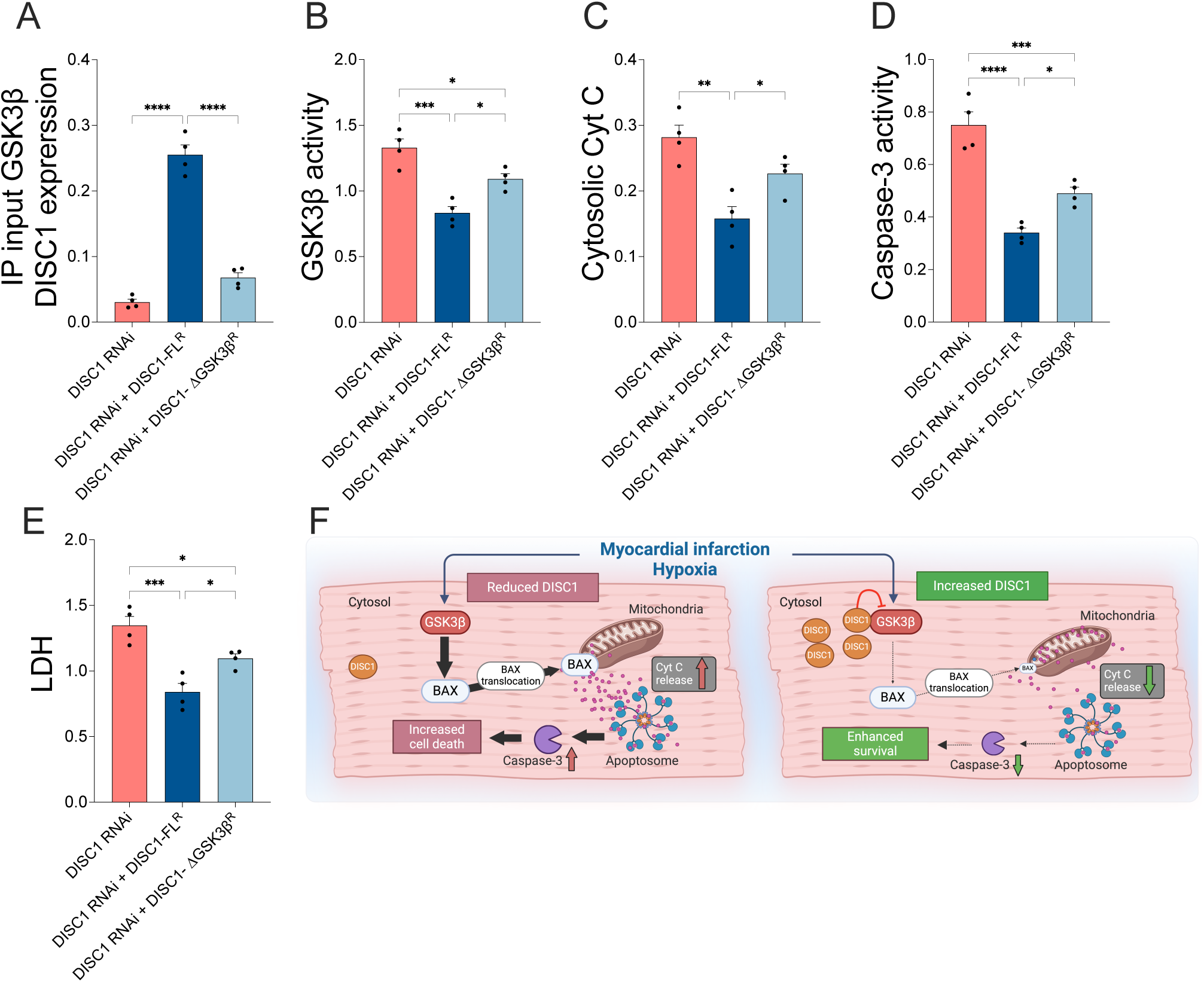
Direct DISC1-GSK3β interaction hinder GSK3β activation and reduces cytosolic Cytochrome C (Cyt C) release and caspase-3 activity. To further asses the role of DISC1 on GSK3β activity during hypoxia, the experiments were performed using the following scenarios; 1) **DISC1 RNAi**, expression of the not resistant to shRNA full length (FL) (DISC1-FL) + co-treatment by shRNA to silence the DISC1, leaving cells with low DISC1 expression as product; 2) **DISC1 RNAi + DISC1-FL^R^**: overexpression of DISC1-FL^R^ that is resistant to shRNA and 3) **DISC1 RNAi + DISC1-ΔGSK3β^R^**, which is the DISC1 overexpression vector with triple mutation at the binding site of GSK3β and resistant to DISC1 RNAi. All experimental groups received treatment with shRNA DISC1 to remove contribution from the endogenous DISC1, thus leaving mainly the product from the respective overexpression vectors, ^R^ denotes resistant to this shRNA (i.e. DISC1-FL^R^ and DISC1-ΔGSK3β^R^). **A**, DISC1-GSK3β interaction by IP; **B**, GSK3β activity; **C**, cytosolic Cyt C levels; **D**, Caspase-3 activity; **E**, Lactate dehydrogenase (LDH) from cardiomyocytes released to medium. **F**, Simplified overview displaying the effect of low DISC1 levels with consequences of interrupted DISC1-GSK3β interaction causing increased cell death, whereas increased DISC1 levels causes maintained DISC1-GSK3β interaction with consequences of reduced GSK3β activity and enhanced survival. All data are presented as mean ± s.e.m. Statistical analyses were performed by one-way ANOVA with Tukey’s post hoc correction for multiple comparisons. *\* P<0.05; **P<0.01; ***P<0.001; ****P<0.0001*.

To further assess the role of DISC1 on GSK3β activity during hypoxia, we repeated the experiments using the DISC1-FL^R^ and compared this to GSK3β activity using the DISC1-ΔGSK3β^R^. We found that the GSK3β activity was increased in presence of DISC1 shRNA with wild type full length DISC1 that is not resistant to shRNA (DISC1 RNAi); GSK3β activity was significantly reduced by overexpression of the DISC1-FL^R^ vector (DISC1 RNAi + DISC1-FL^R^). We also found a modest reduction in GSK3β activity using the DISC1-ΔGSK3β^R^ vector (DISC1 RNAi + DISC1-ΔGSK3β^R^), but most importantly this effect was significantly less than the findings from DISC1-FL^R^. Collectively, these data display that a direct interaction between DISC1 and GSK3β is a major component of the mechanism(s) underlying reduced GSK3β activity by overexpression of DISC1 (**Fig. 4B**).

To further determine the downstream effects of the GSK3β-DISC1 interaction, we used the same experimental design and performed CytC and caspase-3 activity assay. CytC release to cytosol was high when DISC1 was silenced by shRNA DISC1 (DISC1 RNAi), which was rescued by overexpression of DISC1 by DISC1-FL^R^ vector resistant to shRNA (DISC1 RNAi + DISC1-FL^R^), but not by DISC1-ΔGSK3β^R^ (DISC1 RNAi + DISC1-ΔGSK3β^R^) (**Fig. 4C**). These data were also confirmed for the Caspase-3 activity (**Fig. 4D**). LDH released from cardiomyocytes to the medium as a marker for cell death confirmed the same results (**Fig. 4E**). The collective evidence from these studies unequivocally establishes the crucial role of the direct interaction between DISC1 and GSK3β in conferring protection against cell death induced by hypoxic conditions. This interaction in cell death signaling processes is pivotal in maintaining the resilience of cardiomyocytes under hypoxic stress. The disruption of this interaction significantly compromises the resistance of cardiomyocytes to hypoxia, rendering them highly susceptible to cell death.

## Discussion

Cardioprotective therapies targeting reperfusion-related injury remain among the top 10 unmet clinical needs in cardiology.^4–6,33^ This study addresses these existing gaps and sheds light on a discovery that may hold significant implications for addressing key components of ischemic damage on the heart. The unbiased identification of DISC1 in an *in vitro* cardiomyocyte model of ischemia, as unveiled by RNA sequencing, sparked scientific interest in the potential role of DISC1 as a novel regulator of signaling pathways integral for ischemic cardioprotection. This discovery was further substantiated by GWAS enrichment analyses demonstrating an overrepresentation of risk gene products for cardiologic diseases among DISC1 protein interactors.

In this study, we present evidence of a major detrimental effect of reduced DISC1 protein levels on myocardial infarct size in mouse hearts, correlation between low expression levels of DISC1 and increased cTnT release that indicate more severe ischemia-reperfusion injury following surgery in CABG patients, as well as *in vitro* cardiomyocyte survival. Increased ischemic damage is due to the disruption of signaling pathways that are critical for cardioprotection. Although DISC1 is known to interact with many proteins as an intracellular hub, we have pinpointed that its specific interaction with GSK3β is essential for cardioprotection. Specifically, we show that reduced DISC1 levels in cardiomyocytes limits the ability for DISC1 to bind directly to GSK3β and silence the downstream cell death signalling induced by GSK3β activation. In stark contrast, the augmentation of DISC1 protein levels resulted in enhanced cardiomyocyte survival and the activation of cardioprotective signaling via via maintenance of direct DISC1-GSK3β interaction which inhibited the GSK3β activity.

At the mechanistic level, our data on increased cardiomyocyte injury following ischemic and hypoxic challenges by DISC1 depletion are explained by activation of apoptotic signaling trough GSK3β activation. Here, we reported that knockdown of DISC1 enhanced the activation of GSK3β, whereas overexpression of DISC1 reduced GSK3β activity through direct interaction. These data are important, as recent studies have demonstrated that GSK3β inhibition contributes to cardiac protection in both preconditioning ^34^ and postconditioning.^35,36^ This protection due to GSK3β inhibition has been linked to hindering opening of the mPTP ^34,36–38^ and reduced BAX integration in the mitochondrial membrane and MAC pores formation,^39,40^ consequently reducing the release of proapoptotic molecules such as CytC and activation of caspase-3, which is the key regulatory event that induces terminal effector events in the apoptotic cascade.^41–44^ ^45^ All these effects were verified in the present study. Our investigation illuminates the pivotal role of DISC1 as a central conductor within the myocardium, coordinating a network of intracellular processes that collectively confer protection against ischemic heart disease. The importance of maintaining elevated levels of DISC1 for cardioprotection against ischemia-induced cell death is underscored, particularly in light of its direct interaction with GSK3β.

The biphasic expression of DISC1 following ischemic/hypoxic stress is noteworthy. Our data indicate that during the initial phase, DISC1 expression is elevated, suggesting the activation of protective compensatory mechanisms. However, under conditions of chronic, persistent cellular hypoxia, DISC1 expression diminishes. Maintaining DISC1 levels, *i.e.*, the ability to resist this reduction during ischemia/cellular hypoxia, appears crucial for survival. At the molecular level, DISC1 functions as a shield, mitigating the harmful effects of hypoxia on cardiomyocytes. When this shield is compromised or lost, cardiomyocytes become vulnerable, resulting in an increased rate of cell death. This vulnerability leads to a higher incidence of cell death and more extensive myocardial infarction following cardiac ischemia which also were displayed trough the correlation between low DISC1 RNA expression in blood and increased release of cTnT following surgery in CABG patients.

Given that DISC1 protein stability may be enhanced by Phosphodiesterase 4B (PDE4B)^22^, and considering that PDE4 is a prominent target in drug discovery and development, medications targeting PDE4 could potentially be applied or repurposed in this context. The mechanism of DISC1 downregulation remains unclear, and to date, there have been no significant efforts to elucidate the transcriptional mechanisms regulating DISC1. Investigating the transactivation and stability of DISC1 mRNA could represent a novel approach for drug discovery. Additionally, the role of the 26 specific overexpressed miRs in regulating DISC1 protein stability warrants further exploration. These miRs have been identified as key regulators that may influence DISC1 levels by targeting its mRNA, thereby affecting its stability and expression. The role of miRs as regulatory factors of the DISC1 interactome has also been previously addressed in neuronal cells.^21^ Studying these miRs could provide valuable insights into maintaining DISC1 levels and developing novel therapeutic strategies for cardioprotection. Consequently, understanding the transcriptional and post-transcriptional regulation of DISC1 is crucial for developing interventions aimed at preserving its protective functions in both neuronal and cardiac contexts.

## Materials and methods

### RNA sequencing data - Library construction and sequencing

RNeasy Mini Kit from Qiagen (Qiagen Norge, Oslo, Norway) was used for RNA extraction according to the manufacturer’s instructions. RNA concentration was measured using a Qubit® RNA HS Assay Kit on a Qubit® 3.0 Fluorometer (Thermo Fisher Scientific Inc., Waltham, MA, USA). Integrity was assessed using an Agilent RNA 6000 Pico Kit on a 2100 Bioanalyzer instrument (Agilent Technologies, Santa Clara, CA, USA).

RNA sequencing libraries were prepared using a TruSeq Stranded mRNA kit (Illumina, San Diego, CA, USA) according to the manufacturer’s instructions. In brief, 300 ng of total RNA was used as the starting material. First, mRNA was purified from the total RNA using poly-T oligo-attached magnetic beads, followed by random fragmentation using divalent cations at 94°C for 4 min. First- and second strand cDNAs were synthesized using random oligonucleotides and SuperScript II, followed by DNA polymerase I and RNase H. Exonuclease/polymerase was used to produce blunted overhangs. Illumina dual index adapter oligonucleotides were ligated to cDNA after 3’ end adenylation. DNA fragments were enriched by 15 cycles of PCR. The libraries were purified using AMPure XP (Beckman Coulter, Inc., Indianapolis, IN, USA), quantitated by qPCR using the KAPA Library Quantification Kit (Kapa Biosystems, Inc., Wilmington, MA, USA) and validated using the Agilent High Sensitivity DNA Kit on a Bioanalyzer (Agilent Technologies, Santa Clara, CA, USA). The size range of the DNA fragments was measured to be in the range of app. 200-1000 bp and peaked around 285 bp.

Libraries were normalized and pooled to 2.7 pM and subjected to clustering on two NextSeq 500 high output flow cells (Illumina, San Diego, CA, USA). Finally, single-read sequencing was performed for 75 cycles on a NextSeq 500 instrument (Illumina, Inc. San Diego, CA, USA) according to the manufacturer’s instructions. Base calling was performed on a NextSeq 500 instrument by RTA 2.4.6. FASTQ files were generated using bcl2fastq2 Conversion Software v2.20.0.422 (Illumina, Inc. San Diego, CA, USA). FASTQ files were filtered and trimmed (fastp v0.20.0), and transcript counts were generated using quasi alignment (Salmon v1.3.0) to the transcriptome reference sequence (Ensembl, GRCh38 release 103).

Transcript sequences were imported into R statistical software and aggregated to gene counts using the tximport (v1.14.0) bioconductor package. Gene counts were normalized and analyzed for differential expression using the DESeq2 bioconductor package. DESeq2 is a specialized software for the analysis of RNA-seq data that builds a generalized linear model under the assumption of negative binomial distributed values and uses the Wald statistic for significance testing.

### Regulatory networks in hypoxia-stimulated hiPSC-CM relationships between upregulated miRs and their downregulated mRNA targets

Establishing functional relationships between those differentially expressed RNAs, we generated regulatory networks that depicted negative correlations between overexpressed miRNAs and their putative downregulated mRNA transcripts. Regulatory networks in hypoxia-stimulated hiPSC-CM were displayed by the predicted functional relationships between upregulated miRs and their downregulated mRNA targets. All predictions were made by using the MiRWalk 2.0 algorithm.^46^ Protein-protein interactions were extracted from the STRING database.^47^ ***Supporting material for analyses included in separate Excel file, Appendix 3.***

### Cardiac patient biopsies and blood samples

Biopsies were obtained from patients scheduled for elective CABG surgery at St. Olavs Hospital, Trondheim, Norway (stored in local biobank: ID275364). The patients were post-MI <15 days prior to CABG surgery (MI, N=5) versus patients with no previously defined MI (control, N=4). Biopsies were taken before aortic cross-clamping from nonfibrotic and viable left ventricle mid-myocardium in the remote noninfarcted area using a BioPince™ needle from Angiotech Pharmaceuticals, Inc. (Vancouver, Canada) and snap-frozen in liquid nitrogen. All perioperative procedures were performed according to standard routines of the department. Approved by the Regional Committee for Medical and Health Research Ethics (REK), Norway (Study Id numbers 285936).

Blood samples were taken from locally stored and approved (REK ID 275364) biobanked samples from patients undergoing CABG surgery at St. Olavs Hopsital, Trondheim Norway (N=29) (patients characteristics are presented in ***Supplements Table S2***. The specific project testing the hypothesis that low DISC1 RNA expression in blood causes more cardiac ischemia-reperfusion injury during CABG surgery, measured by increased cardiac Troponin T (cTnT) levels, was approved by REK, Norway (Study Id numbers 560077). High sensitive cTnT was analysed from blood serum samples by immunoassay (Elecsys Troponin T hs, Instrument Roche cobas 8000) at the at the Department of Medical Biochemistry at St. Olavs Hospital, Trondheim, Norway that is accredited according to the NS-EN ISO 15189 laboratory standard. DISC1 RNA in blood was analysed from whole blood samples stored at 80°C freezer in PAXgene Blood RNA Tube (PreAnalytiX,Qiagen, Hilden, Germany).

Blood samples of DISC1 RNA was analysed from the timepoint pre surgery. cTnT blood samples was analysed as delta difference from pre surgery to 24 hours post surgery.

### Animal studies

The mouse model was a DISC1 locus impairment (DISC1-Li) mouse model that is based on a C57BL/6 background with a locus deletion that included the *Tsnax*/*Trax* gene located at 5ʹ to the *Disc1* gene was included to the study. The 40-kb locus deleted covers exons 1, 1b, 2 and 3 and depletes *Tsnax*/*Trax-Disc1* intergenic splicing. miRNA in intron 1, which could alter the functions and levels of DISC1 by targeting at least some DISC1-binding partner genes (such as 14-3-3 and CRMP1/2), was also deleted, as previously described.^8^ (Approved by the Norwegian Animal Research Authority (NARA), Id:13651). Experiments using post myocardial infarction in rats were performed by permanent occlusion of the left anterior descending (LAD) coronary artery, as previously described.^48^ Sham surgery was performed without ligation. Following MI, the animals remained in their cages before they were killed after 4 weeks. (NARA, Id: 9768).

### Langendorff perfused hearts

Global ischemia was performed *ex vivo* on isolated mouse hearts using Langendorff perfusion. The laboratory perfusionist was blinded to the animals included. All animals experiments were performed in randomized order. After removal of the heart, the aorta was cannulated, and the heart was perfused with Krebs-Hensleit solution with a stable aortic pressure of 60 mmHg and immersed in a bath. Solutions were bubbled with 95% O2/5% CO2 and maintained at 37°C. After 20 mins of perfusion, the hearts were subjected to 30 mins of ischemia by stopping the perfusion flow. Hearts were then reperfused for 1 h 40 m before being taken down and frozen at -20°C overnight. The following day, hearts were cut into 1.5 mm transverse sections using a heart slicing matrix and incubated for 10 minutes in 1% 2,3,5-triphenyltetrazolium chloride (TTC) shaking at 37°C. Slices were placed in PBS between glass slides and scanned at 3200 dpi using a flatbed scanner. The second and third slices closest to the apex were used for analysis. Intensity thresholds were determined by a blinded third party to discriminate between black (background), red (viable tissue) and white (nonviable tissue). The percentage of nonviable tissue in a heart was measured by calculating the percentage of white pixels as a total of all non-background pixels across the two section scans. All image analyses were performed using ImageJ. (NARA id: 13651). Exlusion criterias to the study was unsuccesfull Langendorff perfusion and hearts that where not able to mount on the perfusion system.

### Cell culture and treatments

Human AC-16 cardiomyocyte cells (EMD Millipore / Merck Millipore / Merck Life Sciences, Catalogue # SCC109, Darmstadt, Germany, RRID:CVCL_4U18) were cultured in Dulbecco’s modified Eagle’s medium (DMEM): Ham’s F12 (1:1; v/v) with 2 mM Glutamine, 12.5% fetal bovine serum (FBS), and 1% antibiotic/antimycotic mix, Following the established guidelines, procedures, and protocols set by the commercial vendor. To conduct gain-of-function and loss-of-function experiments, AC-16 cells were subjected to reverse transfection in suspension using the DISC1 overexpression vector (Full length DISC1 isoform L ((https://www.uniprot.org/uniprotkb/Q9NRI5/entry#Q9NRI5-1; Transcript: Human NM_018662.3) pRK5-DISC1 overexpression) or the vector encoding *DISC1* shRNA (TRC Lentiviral Human DISC1 shRNA, Dharmacon / Horizon Scientific, Cambridge, UK, Catalogue # RHS3979-201826869, Clone Id: TRCN0000118997; mature antisense sequence TTAATGATCTAATCTCCTCGG).

For experiments to determine the direct association of DISC1-GSK3 interaction we used the pRK5 vector systems with the following modification: Based on domain of DISC1 that is crucial for binding to GSK3(193−236) from the full-length protein sequence and the most potent GSK3β binding region of DISC1^32^ we created a triple mutant where Phenylalanines (F) was replaced by Alanine (A) at the sites. Using the pRK5 myc-hDISC1 plasmid as a template, we introduced the F206A, F210A, and F212A mutations into the hDISC1 sequence. Subsequently, we generated an escape mutant resistant to shRNA (^R^) on the pRK5 myc-hDISC1 full length (DISC1-FL) and the pRK5 myc-hDISC1 F206A, F210A, F212A constructs by mutating the binding site sequence of the DISC1 shRNA while preserving the original amino acid sequence of hDISC1 (Name used in the paper for constructs: **DISC1-ΔGSK3β^R^** and **DISC1-FL^R^**. This approach would avoid an effect of GSK3β interacting with the endogenous DISC1 and at the same time is not able to silence either the the overexpressed **DISC1-FL^R^** or the **DISC1-ΔGSK3β^R^**. The escape mutant was designed based on the same vector encoding *DISC1* shRNA; antisense sequence TTAATGATCTAATCTCCTCGG obtained from, RHS3979-201826869, Clone Id: TRCN0000118997. The complete sequences of the relevant constructs produced in this paper are provided in ***supporting data Appendix 4.***

To overexpress or knock-down *DISC1* expression, AC-16 cells were transfected in suspension *(reverse transfection*) with the respective vectors, using Polyfect® (Qiagen Norge, Oslo, Norway, Catalogue # 301107) in accordance to the manufacturer’s guidelines and standardized procedures ^49,50^. The plasmid load to be transfected was standardized to 1 μg per 0.6 x 10^6^ cells and *scaled-up* or *scaled-down* in accordance to the stipulations of the experimental paradigm ^49,50^. The hypoxia challenge (1% O_2_, 5% CO_2_, and 94% N_2_) was effectuated by incubating the transfected AC-16 cells with the specific *hypoxia medium* (***Supplementary Table S3***). The hypoxia challenge was induced and maintained for the designated duration using the *New Brunswick™ Galaxy® 48 R CO_2_ incubator* (Eppendorf Norge AS, Oslo, Norway). For RNA sequencing of hIPS-CMs vials of 1 × 10^6^ Cor.4U cardiomyocytes were seeded in gelatin pre-coated 48-well plates and maintained in Cor.4U Complete Culture Medium total RNA was extracted from hIPS-cardiomyocytes (Cor.4U cardiomyocytes (Ncardia, Cologne, Germany), that was treated by either by 48 hours in hypoxia or normoxic control conditions at the temperature of at 37°C using the same procedure as for AC16 cardiomyocytes. Cells were thereafter harvested and immediately snap frozen and stored at -80°C freezer for subsequent RNA extraction.

### DISC1 ELISA

The DISC1 protein levels in experimental lysates were measured by using a *sandwich* ELISA immunoassay approach. Briefly, 10 ng of DISC1 *capture* antibody (***Supplementary Table S4***) was immobilized in each well of a 96-well microplate. The experimental cell lysates (equivalent to 50 μg of protein content) were incubated with the immobilized DISC1 *capture* antibody, overnight at 4^0^C. The conditioned cell lysate was discarded, the 96-well microplate wells were washed 3x (15 minutes each) with TBS-T (Tris-buffered saline with 0.1% v/v Tween-20) and incubated with the DISC1 *detection* antibody (***Supplementary Table S4***), overnight at 4^0^C. The 96-well microplate wells were washed 3x (15 minutes each) with TBS-T followed by immunodetection with the HRP (horseradish peroxidase)-conjugated secondary antibody using the HRP-substrate TMB (3,3’,5,5’-Tetramethylbenzidine) (Thermo Fisher Scientific, Oslo, Norway, Catalogue # N301) as a chromophore for the colorimetric read-out (λ_450_). The antibody and signal specificity were established by performing *peptide blocking assay* in the entire gamut of experimental lysates. The DISC1 antibody blocking peptide corresponding to the specific epitope for the DISC1 *detection* antibody (***Supplementary Table S4***) was used for the *peptide blocking assay*. The respective optical density (O.D) values from the *peptide blocking assay* were used for experimental blank correction. Data is expressed as experimental blank corrected O.D_450_ (λ_450_) values from three technical replicates for each of the four biological replicates belonging to each experimental group (n = 4).

### Lactate Dehydrogenase (LDH) Assay

Lactate Dehydrogenase (LDH) levels in the conditioned media were assessed as an indicator of general cell death. The measurement of LDH levels in the conditioned medium was performed using a sandwich ELISA immunoassay technique as previously described.^40^ 20 ng of LDH capture antibody (see ***Supplementary Table S4***) was attached to each well of a 96-well microplate. Experimental sample conditioned media (50 μL) were then incubated with the immobilized LDH capture antibody overnight at 4°C. The conditioned media was removed, and the wells of the 96-well microplate were washed three times (15 minutes each) with TBS-T (Tris-buffered saline with 0.1% v/v Tween-20). The wells were then incubated overnight at 4°C with LDH-A and LDH-B detection antibodies (***Supplementary Table S4***). After incubation, the wells were washed again three times (15 minutes each) with TBS-T, followed by immunodetection using an HRP-conjugated secondary antibody. The HRP-substrate OPD (o-phenylenediamine dihydrochloride) (Thermo Fisher Scientific, Oslo, Norway, Catalogue # 34005) was used as a chromophore for the colorimetric read-out at 450 nm (λ450). Antibody and signal specificity were confirmed through peptide blocking assays across all experimental lysates. The LDH-A and LDH-B antibody blocking peptides (***Supplementary Table S4***) were used for these assays. Optical density (O.D) values from the peptide blocking assays were utilized for experimental blank correction. Data is presented as experimental blank corrected O.D450 (λ450) values from three technical replicates for each of the four biological replicates in each experimental group (n = 4).

### Cellular fractionation to segregate the cytosolic and mitochondrial compartments

Cellular fractionation to obtain the cytosolic and mitochondrial fractions was carried out using the “Mitochondria/Cytosol Fractionation Kit” from Abcam (Catalogue # ab65320, Abcam, Cambridge, UK), following the manufacturer’s instructions as previously described.^40^ AC-16 cells were terminally sub-cultured and plated in 150 mm cell-culture plates to reach the desired confluence (2 x 10^7 cells per plate). After undergoing the specified transfection and experimental procedures, the cells were trypsinized and pelleted by centrifugation at 1000 x g for 5 minutes at 4°C. The cell pellets were then resuspended in 1x Cytosol Extraction Buffer Mix containing DTT (dithiothreitol) and protease inhibitors (provided with the kit) and incubated on ice for 10 minutes. The cells were homogenized on ice using a Dounce homogenizer with 30-50 strokes. The resulting cell lysate was transferred to a 1.5-mL microcentrifuge tube and centrifuged at 700 x g for 10 minutes at 4°C. The supernatant was collected (discarding the pellet) and further centrifuged at 10000 x g for 30 minutes at 4°C. The supernatant obtained constituted the cytosolic fraction, while the pellet represented the mitochondrial fraction. The mitochondrial pellet was either resuspended in 100 μL of 1x PBS (pH 7.4) to obtain intact mitochondria or in 100 μL of Mitochondrial Extraction Buffer Mix (with DTT and protease inhibitors) to obtain the total mitochondrial fraction. Both the cytosolic and mitochondrial fractions were then processed for sandwich ELISA immunoassays to verify their integrity. The mitochondrial fraction’s integrity was confirmed by the presence of COX4 (Cytochrome C Oxidase Subunit 4) and the absence of β-Actin, while the cytosolic fraction’s integrity was validated by the absence of COX4 and the presence of β-Actin.

### Cytochrome C Release Assay

The movement of Cytochrome C from the inter-mitochondrial space to the cytosol was assessed by comparing its relative abundance in the cellular and mitochondrial fractions. The levels of Cytochrome C in these fractions were quantified using a sandwich ELISA immunoassay method as previously outlined.^40^ Briefly, 20 ng of the Cytochrome C *capture* antibody (***Supplementary Table S4***) was immobilized in each well of a 96-well microplate. The respective cytosolic fractions (equivalent to 50 μg of protein content) and mitochondrial fractions (equivalent to 20 μg of protein content) were incubated with the immobilized Cytochrome C *capture* antibody, overnight at 4^0^C. The conditioned cytosolic fractions and mitochondrial fractions were discarded and the 96-well microplate wells were washed 3x (15 minutes each) with TBS-T and incubated with the Cytochrome C *detection* antibody (***Supplementary Table S4***), overnight at 4^0^C. The 96-well microplate wells were washed 3x (15 minutes each) with TBS-T followed by immunodetection with the HRP-conjugated secondary antibody, using the HRP-substrate OPD, as a chromophore for the colorimetric read-out (λ_450_). The antibody and signal specificity were established by performing *peptide blocking assay* in the entire gamut of experimental lysates. The Cytochrome C antibody blocking peptide (Cell Signaling Technology, Danvers, MA, USA, Catalogue # 1033) corresponding to the specific epitope for the Cytochrome C antibody (***Supplementary Table S4***) was used for the *peptide blocking assay*. The respective absorbances from the *peptide blocking assay* were used for experimental blank correction. Data is expressed as experimental blank corrected O.D_450_ (λ_450_) values from three technical replicates for each of the four biological replicates belonging to each experimental group (n = 4).

### Caspase-3 Activity Assay

The enzymatic activity of caspase-3 was assessed spectrophotometrically using the specific substrate Ac-DEVD-p-NA (N-Acetyl-Asp-Glu-Val-Asp-p-nitroanilide) (Sigma Aldrich, Oslo, Norway, Catalogue # 235400-5MG) as previusoly described.^40^ Caspase-3 activity was inferred from the amount of chromophore released due to the proteolysis of Ac-DEVD-p-NA by caspase-3. Briefly, AC-16 cells were terminally sub-cultured and plated in 100 mm cell-culture plates to reach the desired confluence (4 x 10^6 cells per plate). After undergoing the specified transfection and experimental procedures, the cells were trypsinized and pelleted by centrifugation at 1000 x g for 5 minutes. The cell pellets were resuspended in cell lysis buffer (50 mM HEPES, 5 mM CHAPS, pH 7.4) and incubated on ice for 10 minutes to lyse the cells, followed by centrifugation at 12000 x g for 15 minutes to pellet the cell debris. The supernatant containing non-denatured cell lysates (without protease inhibitors) was used for caspase-3 activity determination. The protein concentration of the non-denatured cell lysates was adjusted to 4 µg/µL, and the volume was brought to 180 µL with assay buffer (20 mM HEPES, 1.62 mM CHAPS, 10 mM NaCl, 2 mM EDTA, pH 7.4). For the caspase-3 activity assay, 200 µg of protein (50 µL of lysate at 4 µg/µL diluted to 180 µL) per well of a 96-well microplate was used from each experimental sample. The samples were incubated with the caspase-3 substrate, Ac-DEVD-p-NA (200 µM, 20 µL of 2 mM in a 200 µL assay volume), for 4 hours at 37°C. Caspase-3 activity assays were also conducted in the presence of the caspase-3 inhibitor, Ac-DEVD-CHO (Sigma Aldrich / Merck Millipore / Merck Life Science, Darmstadt, Germany, Catalogue # 235420), to confirm assay specificity and serve as an experimental blank. Absorbance at 405 nm (λ405), corresponding to the amount of chromophore produced as a measure of caspase-3 activity, was determined using a microplate reader. Absorbance values from caspase-3 inhibitor-treated lysates were used for experimental blank correction. Data is presented as experimental blank corrected O.D405 (λ405) values from three technical replicates for each of the four biological replicates in each experimental group (n = 4).

### GSK3β kinase activity assay

GSK3β kinase activity in non-denatured experimental lysates was assessed using an Indirect ELISA immunoassay. The phosphorylation level of the serine residue in the synthetic peptide (RRRPASVPPSPSLSRHS(pS)HQRR), which corresponds to the GSK3β consensus recognition motif in the endogenous substrate muscle glycogen synthase 1 ((pS) indicates phosphorylated-serine), served as a surrogate marker for GSK3β kinase activity in the experimental lysates. 96-well microplates were prepared using Streptavidin-Biotin chemistry. Initially, the wells were coated with 20 pmoles (1.1 μg) of Streptavidin (110 μL of 10 μg/mL Streptavidin in 20 mM potassium phosphate, pH 6.5). This was followed by the immobilization of the biotinylated-GSK3β substrate (GSM [GSK3 substrate peptide], Sigma Aldrich / Merck Life Science, Oslo, Norway, Catalogue # 12-533). The N-terminus biotinylation of the GSK3β substrate-peptide was performed using the “EZ-Link™ NHS-LC-Biotin” biotinylation kit from Thermo Fisher Scientific (Thermo Fisher Scientific, Oslo, Norway, Catalogue # 21336) according to the manufacturer’s guidelines. Briefly, 50 ng of the biotinylated GSK3β substrate-peptide was immobilized in each well of the Streptavidin-coated 96-well microplate. Experimental cell lysates (equivalent to 50 μg of protein) were incubated with the immobilized biotinylated-GSK3β substrate overnight at 4°C, followed by a 24-hour incubation with the specific detection-primary antibody (100 ng/well, 50 μL of 2 μg/mL) directed against the phosphorylated-serine residue (***Supplementary Table S4***). The phosphorylated-serine residue in the biotinylated-GSK3β substrate was then immunodetected using an AP (alkaline phosphatase)-conjugated secondary antibody and the AP-substrate PNPP (p-Nitrophenyl Phosphate, disodium salt) (Thermo Fisher Scientific, Oslo, Norway, Catalogue # 37621) as a chromophore for the colorimetric read-out at 405 nm (λ405). GSK3β kinase activity assays were also conducted in the presence of the GSK3β kinase inhibitor SB-216763 (Sigma Aldrich / Merck Millipore / Merck Life Science, Darmstadt, Germany, Catalogue # S3442) to confirm assay specificity and serve as an experimental blank. Optical density (O.D) values from the GSK3β kinase inhibitor-treated lysates were used for experimental blank correction. Data is presented as experimental blank corrected O.D405 (λ405) values from three technical replicates for each of the four biological replicates in each experimental group (n = 4).

### Quantitative measurement of BAX and BAK in cytosolic and mitochondrial fractions using sandwich ELISA

The levels of BAX and BAK in mitochondrial fractions, cytosolic fractions, and whole-cell lysates were determined using a sandwich ELISA immunoassay. Briefly, 10-30 ng of the respective capture antibodies (***Supplementary Table S4***) were immobilized in each well of 96-well microplates. Mitochondrial fractions (equivalent to 20 μg of protein) and cytosolic fractions (equivalent to 40 μg of protein) were incubated with the immobilized capture antibodies overnight at 4°C. After incubation, the conditioned mitochondrial and cytosolic fractions were discarded, and the wells were washed three times (15 minutes each) with TBS-T. The wells were then incubated overnight at 4°C with the respective detection antibodies (***Supplementary Table S4***). Following this, the wells were washed three times (15 minutes each) with TBS-T and subjected to immunodetection using an HRP-conjugated secondary antibody and the HRP-substrate OPD for colorimetric read-out at 450 nm (λ450). Antibody signal specificity was confirmed through peptide blocking assays across all mitochondrial and cytosolic fractions from experimental cells. The antibody blocking peptides corresponding to the specific epitopes for the detection antibodies used are listed in ***Supplementary Table S4***. Absorbance values from the peptide blocking assays were used for experimental blank correction. Data is presented as experimental blank corrected O.D450 (λ450) values normalized to fold-change, derived from three technical replicates for each of the four biological replicates in each experimental group (n = 4).

### Isolation of heavy mitochondrial fractions for ELISA immunoassays

Heavy mitochondrial fractions were isolated using the “Mitochondria Isolation Kit” from Sigma Aldrich / Merck Millipore (Merck Life Science, Darmstadt, Germany, Catalogue # MITOISO2) as previously described.^40^ Briefly, AC-16 cells were terminally sub-cultured and plated in 150 mm cell-culture plates to reach the desired confluence (2 x 10^7^ cells per plate). After undergoing the specified transfection and experimental procedures, the cells were trypsinized and pelleted by centrifugation at 600 x g for 5 minutes at 4°C. The cell pellets were resuspended in 1.2 mL of lysis buffer (provided with the kit) containing protease inhibitors (also provided with the kit) and incubated on ice for 5 minutes to lyse the cells. This was followed by the addition of twice the volume of extraction buffer (provided with the kit) and centrifugation at 1000 x g for 10 minutes at 4°C. The resulting supernatant was further centrifuged at 3500 x g for 10 minutes at 4°C to obtain the heavy mitochondrial fraction. For quantitative ELISA immunoassays, the pelleted heavy mitochondrial fraction was resuspended in 200 μL of CelLytic™ M Cell Lysis Reagent (provided with the kit) containing protease inhibitors (1:100 [v/v]).

### Isolation and preparation of alkali-resistant outer mitochondrial membrane (OMM)-inserted and alkali-soluble OMM-tethered protein fractions for the quantitative measurement of OMM-inserted and OMM-tethered BAX and BAK

To determine the abundance of oligomeric OMM-inserted BAX and BAK, the pelleted heavy mitochondrial fraction (as described in Section M.9) was resuspended and incubated in 900 μL of 100 mM sodium carbonate (pH 11.5) at 4°C for 45 minutes, followed by ultracentrifugation at 144000 x g at 4°C for 45 minutes. The resulting pellet contained the alkali-resistant OMM-inserted protein fraction with BAX and BAK oligomers, while the supernatant contained the alkali-soluble OMM-tethered protein fraction with BAX and BAK monomers. The supernatants were neutralized with glacial acetic acid, and the alkali-soluble proteins were precipitated using TCA (trichloroacetic acid), followed by centrifugation at 15000 x g at 4°C for 60 minutes. The pellets from both the alkali-resistant OMM-inserted protein fraction (containing BAX and BAK oligomers) and the alkali-soluble OMM-tethered protein fraction (containing BAX and BAK monomers) were resuspended in denaturing RIPA lysis buffer (50 mM Tris, 150 mM NaCl, 0.5% w/v Sodium Deoxycholate, 0.1% w/v SDS, 1% v/v Triton-X, pH 7.4) supplemented with protease and phosphatase inhibitors (Halt™ Protease and Phosphatase Inhibitor Cocktail 100x, Thermo Fisher Scientific, Oslo, Norway, Catalogue # 78446). These fractions were then processed for ELISA immunoassays to quantify the relative abundance of BAX and BAK and to validate the purity and integrity of the fractions. Validation was performed by analyzing the presence and abundance of TOM40 (an OMM-inserted protein), TIM22 (an IMM-inserted protein), HK2 (an OMM-tethered protein), and SDHA (an IMM-tethered protein).

### DISC1 and GSK3β Co-Immunoprecipitation (Co-IP) Analysis

Co-Immunoprecipitation (Co-IP) assays coupled with tandem ELISA immunoassays were performed to determine the relative abundance of the DISC1 mutants in the GSK3β immunoprecipitates. AC-16 cells (4 × 10^6^) seeded in 100 mm plates, transfected and subjected to the experimental interventions (as enunciated earlier), were homogenized using a *non-denaturing* lysis buffer (20 mM Tris, 137 mM Nacl, 2 mM EDTA, 1% Nonidet P-40, 10% glycerol, pH 7.4) supplemented with protease and phosphatase inhibitors. The experimental cell lysates containing the equivalent to 750 μg of total protein was pre-cleared by incubation with protein A/G coated agarose beads for 30 minutes at 4^0^C to reduce the non-specific binding of proteins to the beads. The equivalent of 750 μg of the pre-cleared lysate was incubated separately, with either 5 μg of GSK3β antibody or 5 μg of the corresponding control rabbit IgG antibody (***Supplementary Table S4***), overnight at 4^0^C. The respective immunocomplexes were captured and immobilized by the addition of protein A/G agarose beads and incubation overnight at 4^0^C. The beads containing the immunocomplexes were washed 3x with the *non-denaturing* lysis buffer followed by centrifugation and discarder of the supernatant. The beads were suspended in *denaturing* RIPA buffer (50 mM Tris, 150mM Nacl, 0.1% SDS, 0.5% sodium deoxycholate, 1% Triton X, pH 7.4) supplemented with protease and phosphatase inhibitors and centrifuged to pellet the beads. The supernatant containing the immunoprecipitated proteins was subjected to tandem ELISA with the designated antibodies for DISC1 (***Supplementary Table S4***).

### Integrity and validation of the fractionated mitochondrial and cytosolic compartments for Cytochrome C release assay and quantitative ELISA of BAX and BAK

The integrity of the mitochondrial fraction was confirmed by the presence of COX4 and the absence of β-Actin, while the cytosolic fraction’s integrity was validated by the absence of COX4 and the presence of β-Actin. Levels of COX4 and β-Actin in mitochondrial fractions, cytosolic fractions, and whole-cell lysates were determined using a sandwich ELISA immunoassay. Briefly, 10-30 ng of the respective capture antibodies (***Supplementary Table S4***) were immobilized in each well of 96-well microplates. Mitochondrial fractions (20 μg of protein) and cytosolic fractions (40 μg of protein) were incubated with the immobilized capture antibodies overnight at 4°C. After incubation, the conditioned fractions were discarded, and the wells were washed three times (15 minutes each) with TBS-T. The wells were then incubated overnight at 4°C with the respective detection antibodies (***Supplementary Table S4***). Following this, the wells were washed three times (15 minutes each) with TBS-T and subjected to immunodetection using an HRP-conjugated secondary antibody and the HRP-substrate Amplex Red (10-acetyl-3,7-dihydroxyphenoxazine) (Thermo Fisher Scientific, Oslo, Norway, Catalogue # A22188) for colorimetric read-out at 570 nm (λ570). Antibody signal specificity was confirmed through peptide blocking assays across all mitochondrial and cytosolic fractions from experimental cells. The antibody blocking peptides corresponding to the specific epitopes for the detection antibodies used are listed in ***Supplementary Table S4***. Absorbance values from the peptide blocking assays were used for experimental blank correction. Data is presented as experimental blank corrected O.D570 (λ570) values normalized to fold-change, derived from three technical replicates for each of the four biological replicates in each experimental group (n = 4).

### Validation of the integrity of the alkali-resistant resistant outer mitochondrial membrane (OMM)-inserted and alkali soluble OMM-tethered protein fractions

The integrity of the OMM-inserted protein fraction was validated by the presence of TOM40 and TIM22 concomitant with the absence of SDHA and HK2. The integrity of the OMM-tethered protein fraction was validated by the absence of TOM40 and TIM22 concomitant with the presence of SDHA and HK2. The levels of SDHA, HK2, TOM40, and TIM22, in the mitochondrial fractions, cytosolic fractions, as well as the whole-cell lysates, were determined by *sandwich* ELISA immunoassay. Briefly, 10-30 ng of the respective *capture* antibodies (***Supplementary Table S4***) were immobilized in each well of the respective 96-well microplates. The mitochondrial fractions (equivalent to 20 μg of protein content) and the cytosolic fractions (equivalent to 40 μg of protein content) were incubated with the respective immobilized *capture* antibodies, overnight at 4^0^C. The conditioned mitochondrial fractions and cytosolic fractions were discarded and the respective 96-well microplate wells were washed 3x (15 minutes each) with TBS-T and incubated with the respective *detection* antibodies (***Supplementary Table S4***), overnight at 4^0^C. The 96-well microplate wells were washed 3x (15 min each) with TBS-T, followed by immunodetection with the HRP-conjugated secondary antibody using the HRP-substrate TMB (3,3ʹ,5,5ʹ-tetramethylbenzidine) (Thermo Fisher Scientific, Oslo, Norway, Catalogue # N301) as a chromophore for the colorimetric read-out (λ_450_).The antibody signal specificity was established by performing peptide blocking assays in the entire gamut of experimental samples. The antibody blocking peptides corresponding to the specific epitopes for the respective *detection* antibodies used are enumerated in ***Supplementary Table S4***. The respective absorbances from the peptide blocking assays were used for the experimental blank correction. Data is expressed as experimental blank corrected O.D_450_ (λ_450_) values normalized to *fold-change*, emanating from three technical replicates for each of the four biological replicates belonging to each experimental group (n = 4).

### Quantitative real-time RT-PCR analysis

Total RNA was extracted from treated AC-16 cardiomyocytes using the “RNeasy Mini Kit” from Qiagen (Qiagen Norge, Oslo, Norway, Catalogue # 74104) following the manufacturer’s instructions and protocols. cDNA was obtained by reverse transcribing 1 μg of extracted total RNA using the “QuantiTect Reverse Transcription Kit” from Qiagen (Qiagen Norge, Oslo, Norway, Catalogue # 205313). cDNA amplification was performed using the “miScript SYBR Green PCR Kit” from Qiagen (Qiagen Norge, Oslo, Norway, Catalogue # 218075) following the manufacturer’s instructions and protocols. The primer pair used to amplify the *DISC1 mRNA transcript variant L* (NCBI Reference Sequence: NM_018662.3) was as follows: *forward primer 5’-AGAGAGAGAAGGGCTGGAGG-3’* and *reverse primer 5’-TGGAGCTGTAGGCTCTGGAT-3’.* The expression of specific amplified *DISC1* transcripts was normalized to the expression of *actin beta (ACTB)* (NCBI Reference Sequence: NM_001101.5). The primer pair used to amplify the *ACTB mRNA transcript* was as follows: *forward primer 5’-CTCGCCTTTGCCGATCC-3’* and *reverse primer 5’-TCTCCATGTCGTCCCAGTTG-3’.* The expression of specific amplified *DISC1* transcripts was normalized to the expression of *ACTB* using the *delta delta C_T_* method.^51^

*<u>Cardiac samples from CABG left ventricle myocardium</u>* biopsies were stored at -80 °C in RNAlater. After thawing, the biopsies were washed with RNase-free water and excess water remover by bloting againg paper and transferring to new tubes with 4 ceramic beads and 300 µl of RTL lysis buffer. RNA extraction was performed using an RNeasy kit for fibrous tissues (Qiazol) according the manufacturer’s protocol. cDNA was obtained using a Quantitech reverse transcription kit (Qiagen) according to the manufacturer’s protocol. Quantification was performed by DISC1 Primer Assays from Qiagen (Qiagen QuantiTect Primer Assay (QT01019060) normalized to B2M as a housekeeping gene. For detection of DISC1 RNA expression in *<u>blood from CABG patients</u>*, RNA was isolated from PAXgene tubes (PreAnalytiX,Qiagen, Hilden, Germany), using NAxtra™ Blood total nucleic acid extraction kit (Lybe Scientific, Trondheim, Norway) according to manufacturer’s protocol.^52^ Briefly, blood was spun down and supernatant removed, and protein digested using Proteinase K. Using a Kingfisher Flex (ThermoFisher), samples were then incubated in a lysis buffer, nucleic acid bound to magnetic beads, DNA was digested, and samples were washed. RNA was quantified using a Qubit RNA HS Assay Kit (Invitrogen). cDNA was reverse transcribed from these samples using QuantiTect Reverse Transcription Kit (Qiagen) according to manufacturer’s protocol. DISC1 mRNA was quantified by RT-PCR, using QuantiTect SYBR Green PCR Kit (Qiagen) according to the manufacturer’s protocol. Reactions were run in a CFX Opus Real Time PCR system (Bio-Rad). The primer pair used to amplify the DISC1 mRNA transcript variant L was the same as described above for human AC16 cardiomyocytes. The expression of specific amplified *DISC1* transcripts was normalized to the expression of Homo sapiens hypoxanthine phosphoribosyltransferase (HPRT) (NCBI Reference Sequence: NM_000194). The primer pair used to amplify the HPRT mRNA transcript was as follows: *forward primer 5’-CCTGGCGTCGTGATTAGTGAT -3’* and *reverse primer 5’-* AGACGTTCAGTCCTGTCCATAA *-3’* (Source: Primerbank 164518913c1^53^). The expression of specific amplified *DISC1* transcripts was normalized to the expression of HPRT using the *delta delta C_T_* method.^51^

*<u>For rat tissue</u>*, cDNA was obtained using the QuantiTect Reverse Transcription Kit (Qiagen) following the manufacturer’s instructions. qRT-PCRs were performed using a QuantiTect SYBR Green PCR Kit (Qiagen). The primer pair used to amplify DISC1 in rats was as follows: *forward primer 5’-GACAGTGGTTGTCGGCAAGAAT-3’* and *reverse primer 5’-TGCTCCAAGCTACATCAAGGC-3’.* PCR amplification was performed using “*CFX96 Real-Time PCR Detection*” from Bio-Rad (Bio-Rad Norway AS, Oslo, Norway) in all samples.

### Declaration of generative AI and AI-assisted technologies in the writing process

During the preparation of this work the authors used [Microsoft Copilot] in order to lighty improve language and readability. After using this tool, the authors reviewed and edited the content as needed and take full responsibility for the content of the publication.

### Statistical analyses

Statistical analysis was performed with GraphPad Prism 10 (GraphPad Software, San Diego, CA, USA). Quantitative data for all the assays are presented as Mean values ± s.e.m. (Standard Error of the mean). Relevant statistical methods are presented in each data set.

## Supporting information

Merged supplementary Figures S1-S6 Data S1. Tables S1-S4

## Acknowledgment

The current project was financed by grants from the Research Council Norway – FRIPRO ID 316002 to MAH. The project was also funded by Foundation Dam (grant number: FOR555614) to MAH. This study is further supported by The National Institute of Mental Health Grants MH-094268, MH-105660, and MH129480 to AS. The RNA library prep, sequencing and bioinformatics analysis were performed in close collaboration with the Genomics Core Facility (GCF), Norwegian University of Science and Technology (NTNU). We especially acknowledge Arnar Flatberg for bioinformatics analyses. GCF is funded by the Faculty of Medicine and Health Sciences at NTNU and Central Norway Regional Health Authority. We acknowledge Dr. Karin Garten for contribution to the method of determining infarct size in TTC stained cardiac tissue.

## Author contributions

Conceptualization: MAH

Methodology: GM, MP, NRS, RER, FJE, KY, KI, CAAC, KP, RKN, MAH

Investigation: GM, MP, KH, AW, NRS, RB, RER, RKN, FJE, KY, KI, CAAC, KP, GB, AS, MAH

Visualization: MAH, GM, KP, NRS

Supervision: MAH, AS

Writing—original draft: MAH

Writing—review & editing: GM, MP, KH, AW, NRS, RB, RER, FJE, KY, KI, CAAC, RKN, KP, GB, AS, MAH

## Notes

### Competing Interest Statement

The authors have declared no competing interest.

### Summary of Updates

The initial part of both manuscripts centers around an unbiased genome-wide screening designed to identify key molecular drivers of ischemic heart conditions. In both versions DISC1 emerged as the top candidate, an unexpected finding given its established role in neurobiology. The earlier version introduced DISC1 as a novel molecule with potential relevance to cardiac biology. The updated version of the manuscript significantly expands on this rationale. It provides a more detailed and structured explanation of how DISC1 was prioritized for mechanistic investigation; this includes refined description on the bioinformatics analyses. The reframing of the manuscript marks a conceptual shift. DISC1 is no longer presented merely as a surprising candidate; it is positioned as a pivotal regulator of cardiac resilience to ischemia and cellular hypoxia. The new manuscript emphasize a more established and mechanistically grounded understanding the role of DISC1 in cardioprotection. Data in the previous version was largely descriptive showing that downregulation and overexpression of DISC1 affected multiple signaling pathways for cardioprotection. Causality was not demonstrated; The updated version addresses this gap by focusing on the DISC1-GSK3β interaction as a central mechanism through a newly introduced set of experiments: The new version update by using a triple mutant construct to disrupt the binding between DISC1 and GSK3β (New figure 4). These experiments demonstrate that the protective effects of DISC1 are contingent upon its ability to bind GSK3β. This direct mechanistic link elevates the scientific rigor of the study and transforms the narrative from observational to causative. The updated manuscript strengthens its translational relevance by incorporating clinical data from coronary artery bypass grafting (CABG) patients. It shows that low DISC1 RNA expression in blood correlates significantly with increased cardiac troponin T release 24 hours post-surgery. This correlation (r = - 0.32, P = 0.045) provides compelling evidence that DISC1 may play a protective role not only in experimental models but also in human cardiac pathology. The updated version revisits key experiments from the original manuscript, including GSK3β activity assays and LDH release under hypoxic stress (New figure 3). These experiments are now more precisely contextualized within the framework of DISC1 interacting with GSK3β. The manuscript as a whole has been refined to deliver a more focused and coherent mechanistic narrative. This version move beyond broad associations to establish DISC1 as a critical molecular shield against ischemic damage in the heart.

