## Supporting Appendix 4 for "Pivotal role of Disrupted-in-Schizophrenia 1 (DISC1) in cardiac resilience to ischemic stress"

Complete sequences of vectors established in this paper:

**Wildtype: pRK5 myc-hDISC1 (myc-DISC1 is highlighted in green)**

GACATTGATTATTGACTAGTTATTAATAGTAATCAATTACGGGGTCATTAGTTCATAGCCCATATATGGAGTTCCGCGTTACATAACTTACGGTAAATGGCCCGCCTGGCTGACCGCCCAACGACCCCCGCCCATTGACGTCAATAATGACGTATGTTCCCATAGTAACGCCAATAGGGACTTTCCATTGACGTCAATGGGTGGAGTATTTACGGTAAACTGCCCACTTGGCAGTACATCAAGTGTATCATATGCCAAGTACGCCCCCTATTGACGTCAATGACGGTAAATGGCCCGCCTGGCATTATGCCCAGTACATGACCTTATGGGACTTTCCTACTTGGCAGTACATCTACGTATTAGTCATCGCTATTACCATGGTGATGCGGTTTTGGCAGTACATCAATGGGCGTGGATAGCGGTTTGACTCACGGGGATTTCCAAGTCTCCACCCCATTGACGTCAATGGGAGTTTGTTTTGGCACCAAAATCAACGGGACTTTCCAAAATGTCGTAACAACTCCGCCCCATTGACGCAAATGGGCGGTAGGCGTGTACGGTGGGAGGTCTATATAAGCAGAGCTCGTTTAGTGAACCGTCAGATCGCCTGGAGACGCCATCCACGCTGTTTTGACCTCCATAGAAGACACCGGGACCGATCCAGCCTCCGCGGCCGGGAACGGTGCATTGGAACGCGGATTCCCCGTGCCAAGAGTGACGTAAGTACCGCCTATAGAGTCTATAGGCCCACCCCCTTGGCTTCGTTAGAACGCGGCTACAATTAATACATAACCTTATGTATCATACACATACGATTTAGGTGACACTATAGAATAACATCCACTTTGCCTTTCTCTCCACAGGTGTCCACTCCCAGGTCCAACTGCACCTCGGTTCTATCGATTGAATTCCCGAACCGACAGTCGGTCTCTTCACCAAGGCCATTCGCGCCACCATGGAGCAAAAGCTCATTTCTGAGGAAGATCTCAATGGTGGTGGTGGTGGGTCGACCATGCCAGGCGGGGGTCCTCAGGGCGCCCCAGCCGCCGCCGGCGGCGGCGGCGTGAGCCACCGCGCAGGCAGCCGGGATTGCTTACCACCTGCAGCGTGCTTTCGGAGGCGGCGGCTGGCACGGAGGCCGGGCTACATGAGAAGCTCGACAGGGCCTGGGATCGGGTTCCTTTCCCCAGCAGTGGGCACACTGTTCCGGTTCCCAGGAGGGGTGTCTGGCGAGGAGTCCCACCACTCGGAGTCCAGGGCCAGACAGTGTGGCCTTGACTCGAGAGGCCTCTTGGTCCGGAGCCCTGTTTCCAAGAGTGCAGCAGCCCCTACTGTGACCTCTGTGAGAGGAACCTCGGCGCACTTTGGGATTCAGCTCAGAGGTGGCACCAGATTGCCTGACAGGCTTAGCTGGCCGTGTGGCCCTGGGAGTGCTGGGTGGCAGCAAGAGTTTGCAGCCATGGATAGTTCTGAGACCCTGGACGCCAGCTGGGAGGCAGCCTGCAGCGATGGAGCAAGGCGTGTCCGGGCAGCAGGCTCTCTGCCATCAGCAGAGTTGAGTAGCAACAGCTGCAGCCCTGGCTGTGGCCCTGAGGTCCCCCCAACCCCTCCTGGCTCTCACAGTGCCTTTACCTCAAGCTTTAGCTTTATTCGGCTCTCGCTTGGCTCTGCCGGGGAACGTGGAGAAGCAGAAGGCTGCCCACCATCCAGAGAGGCTGAGTCCCATTGCCAGAGCCCCCAGGAGATGGGAGCCAAAGCTGCCAGCTTGGACGGGCCTCACGAGGACCCGCGATGTCTCTCTCGGCCCTTCAGTCTCTTGGCTACACGGGTCTCTGCAGACTTGGCCCAGGCCGCAAGGAACAGCTCCAGGCCAGAGCGTGACATGCATTCTTTACCAGACATGGACCCTGGCTCCTCCAGTTCTCTGGATCCCTCACTGGCTGGCTGTGGTGGTGATGGGAGCAGCGGCTCAGGGGATGCCCACTCTTGGGACACCCTGCTCAGGAAATGGGAGCCAGTGCTGCGGGACTGCCTGCTGAGAAACCGGAGGCAGATGGAGGTAATATCCTTAAGATTAAAACTTCAGAAACTTCAGGAAGATGCAGTTGAGAATGATGATTATGATAAAGCTGAGACGTTACAACAAAGATTAGAAGACCTGGAACAAGAGAAAATCAGCCTGCACTTTCAACTTCCTTCAAGGCAGCCAGCTCTTAGCAGTTTCCTGGGTCACCTGGCAGCACAAGTCCAGGCTGCCTTGCGCCGTGGGGCCACTCAGCAGGCCAGCGGAGATGACACCCACACCCCACTGAGAATGGAGCCGAGGCTGTTGGAACCCACTGCTCAGGACAGCTTGCACGTGTCCATCACGAGACGAGACTGGCTTCTTCAGGAAAAGCAGCAGCTACAGAAAGAAATCGAAGCTCTCCAAGCAAGGATGTTTGTGCTGGAAGCCAAAGATCAACAGCTGAGAAGGGAAATAGAGGAGCAAGAGCAGCAACTCCAGTGGCAGGGCTGCGACCTGACCCCACTGGTGGGCCAGCTGTCCCTGGGTCAGCTGCAGGAGGTCAGCAAGGCCTTGCAGGACACCCTGGCCTCAGCCGGTCAGATTCCCTTCCATGCAGAGCCACCGGAAACCATAAGGAGCCTCCAGGAAAGAATAAAATCCCTCAACTTGTCACTTAAAGAAATCACTACTAAGGTGTGTATGAGTGAGAAATTCTGCAGCACCCTGAGGAAGAAAGTTAACGATATTGAAACCCAACTACCAGCCTTGCTTGAAGCCAAAATGCATGCCATATCAGGAAACCATTTCTGGACGGCTAAAGACCTCACCGAGGAGATTAGATCATTAACATCAGAGAGAGAAGGGCTGGAGGGACTCCTCAGCAAGCTGTTGGTGTTGAGTTCCAGGAATGTCAAAAAGCTGGGAAGTGTTAAAGAAGATTACAACAGACTGAGAAGAGAAGTGGAGCACCAGGAGACTGCCTATGAAACAAGTGTGAAGGAAAATACTATGAAGTACATGGAAACACTTAAGAATAAACTGTGCAGCTGCAAGTGTCCACTGCTTGGGAAAGTGTGGGAAGCTGACTTGGAAGCTTGTCGATTGCTTATCCAGAGCCTACAGCTCCAGGAAGCCAGGGGAAGCCTGTCTGTAGAAGATGAGAGGCAGATGGATGACTTAGAGGGAGCTGCTCCTCCTATTCCCCCCAGGCTCCACTCCGAGGATAAAAGGAAGACCCCTTTGAAGGTATTGGAAGAATGGAAGACTCACCTCATCCCCTCTCTGCACTGTGCTGGAGGTGAACAGAAAGAGGAATCTTACATCCTTTCTGCAGAACTTGGAGAAAAGTGTGAAGACATAGGCAAGAAGCTATTGTACTTGGAAGATCAACTTCACACAGCAATCCACAGTCATGATGAAGATCTCATTCAGTCTCTCAGGAGGGAGCTCCAGATGGTGAAGGAAACTCTGCAGGCCATGATCCTGCAGCTCCAGCCAGCAAAGGAGGCGGGAGAAAGAGAAGCTGCAGCTTCCTGCATGACAGCTGGTGTCCACGAAGCACAAGCCTGAGCGGCCGCTAAGTAAGTAAGGATCCCCAGCTTGGCCGCCATGGCCCAACTTGTTTATTGCAGCTTATAATGGTTACAAATAAAGCAATAGCATCACAAATTTCACAAATAAAGCATTTTTTTCACTGCATTCTAGTTGTGGTTTGTCCAAACTCATCAATGTATCTTATCATGTCTGGATCGGGAATTAATTCGGCGCAGCACCATGGCCTGAAATAACCTCTGAAAGAGGAACTTGGTTAGGTACCTTCTGAGGCGGAAAGAACCAGCTGTGGAATGTGTGTCAGTTAGGGTGTGGAAAGTCCCCAGGCTCCCCAGCAGGCAGAAGTATGCAAAGCATGCATCTCAATTAGTCAGCAACCAGGTGTGGAAAGTCCCCAGGCTCCCCAGCAGGCAGAAGTATGCAAAGCATGCATCTCAATTAGTCAGCAACCATAGTCCCGCCCCTAACTCCGCCCATCCCGCCCCTAACTCCGCCCAGTTCCGCCCATTCTCCGCCCCATGGCTGACTAATTTTTTTTATTTATGCAGAGGCCGAGGCCGCCTCGGCCTCTGAGCTATTCCAGAAGTAGTGAGGAGGCTTTTTTGGAGGCCTAGGCTTTTGCAAAAAGCTGTTAACAGCTTGGCACTGGCCGTCGTTTTACAACGTCGTGACTGGGAAAACCCTGGCGTTACCCAACTTAATCGCCTTGCAGCACATCCCCCTTTCGCCAGCTGGCGTAATAGCGAAGAGGCCCGCACCGATCGCCCTTCCCAACAGTTGCGCAGCCTGAATGGCGAATGGCGCCTGATGCGGTATTTTCTCCTTACGCATCTGTGCGGTATTTCACACCGCATACGTCAAAGCAACCATAGTACGCGCCCTGTAGCGGCGCATTAAGCGCGGCGGGTGTGGTGGTTACGCGCAGCGTGACCGCTACACTTGCCAGCGCCCTAGCGCCCGCTCCTTTCGCTTTCTTCCCTTCCTTTCTCGCCACGTTCGCCGGCTTTCCCCGTCAAGCTCTAAATCGGGGGCTCCCTTTAGGGTTCCGATTTAGTGCTTTACGGCACCTCGACCCCAAAAAACTTGATTTGGGTGATGGTTCACGTAGTGGGCCATCGCCCTGATAGACGGTTTTTCGCCCTTTGACGTTGGAGTCCACGTTCTTTAATAGTGGACTCTTGTTCCAAACTGGAACAACACTCAACCCTATCTCGGGCTATTCTTTTGATTTATAAGGGATTTTGCCGATTTCGGCCTATTGGTTAAAAAATGAGCTGATTTAACAAAAATTTAACGCGAATTTTAACAAAATATTAACGTTTACAATTTTATGGTGCACTCTCAGTACAATCTGCTCTGATGCCGCATAGTTAAGCCAGCCCCGACACCCGCCAACACCCGCTGACGCGCCCTGACGGGCTTGTCTGCTCCCGGCATCCGCTTACAGACAAGCTGTGACCGTCTCCGGGAGCTGCATGTGTCAGAGGTTTTCACCGTCATCACCGAAACGCGCGAGACGAAAGGGCCTCGTGATACGCCTATTTTTATAGGTTAATGTCATGATAATAATGGTTTCTTAGACGTCAGGTGGCACTTTTCGGGGAAATGTGCGCGGAACCCCTATTTGTTTATTTTTCTAAATACATTCAAATATGTATCCGCTCATGAGACAATAACCCTGATAAATGCTTCAATAATATTGAAAAAGGAAGAGTATGAGTATTCAACATTTCCGTGTCGCCCTTATTCCCTTTTTTGCGGCATTTTGCCTTCCTGTTTTTGCTCACCCAGAAACGCTGGTGAAAGTAAAAGATGCTGAAGATCAGTTGGGTGCACGAGTGGGTTACATCGAACTGGATCTCAACAGCGGTAAGATCCTTGAGAGTTTTCGCCCCGAAGAACGTTTTCCAATGATGAGCACTTTTAAAGTTCTGCTATGTGGCGCGGTATTATCCCGTATTGACGCCGGGCAAGAGCAACTCGGTCGCCGCATACACTATTCTCAGAATGACTTGGTTGAGTACTCACCAGTCACAGAAAAGCATCTTACGGATGGCATGACAGTAAGAGAATTATGCAGTGCTGCCATAACCATGAGTGATAACACTGCGGCCAACTTACTTCTGACAACGATCGGAGGACCGAAGGAGCTAACCGCTTTTTTGCACAACATGGGGGATCATGTAACTCGCCTTGATCGTTGGGAACCGGAGCTGAATGAAGCCATACCAAACGACGAGCGTGACACCACGATGCCTGTAGCAATGGCAACAACGTTGCGCAAACTATTAACTGGCGAACTACTTACTCTAGCTTCCCGGCAACAATTAATAGACTGGATGGAGGCGGATAAAGTTGCAGGACCACTTCTGCGCTCGGCCCTTCCGGCTGGCTGGTTTATTGCTGATAAATCTGGAGCCGGTGAGCGTGGGTCTCGCGGTATCATTGCAGCACTGGGGCCAGATGGTAAGCCCTCCCGTATCGTAGTTATCTACACGACGGGGAGTCAGGCAACTATGGATGAACGAAATAGACAGATCGCTGAGATAGGTGCCTCACTGATTAAGCATTGGTAACTGTCAGACCAAGTTTACTCATATATACTTTAGATTGATTTAAAACTTCATTTTTAATTTAAAAGGATCTAGGTGAAGATCCTTTTTGATAATCTCATGACCAAAATCCCTTAACGTGAGTTTTCGTTCCACTGAGCGTCAGACCCCGTAGAAAAGATCAAAGGATCTTCTTGAGATCCTTTTTTTCTGCGCGTAATCTGCTGCTTGCAAACAAAAAAACCACCGCTACCAGCGGTGGTTTGTTTGCCGGATCAAGAGCTACCAACTCTTTTTCCGAAGGTAACTGGCTTCAGCAGAGCGCAGATACCAAATACTGTTCTTCTAGTGTAGCCGTAGTTAGGCCACCACTTCAAGAACTCTGTAGCACCGCCTACATACCTCGCTCTGCTAATCCTGTTACCAGTGGCTGCTGCCAGTGGCGATAAGTCGTGTCTTACCGGGTTGGACTCAAGACGATAGTTACCGGATAAGGCGCAGCGGTCGGGCTGAACGGGGGGTTCGTGCACACAGCCCAGCTTGGAGCGAACGACCTACACCGAACTGAGATACCTACAGCGTGAGCTATGAGAAAGCGCCACGCTTCCCGAAGGGAGAAAGGCGGACAGGTATCCGGTAAGCGGCAGGGTCGGAACAGGAGAGCGCACGAGGGAGCTTCCAGGGGGAAACGCCTGGTATCTTTATAGTCCTGTCGGGTTTCGCCACCTCTGACTTGAGCGTCGATTTTTGTGATGCTCGTCAGGGGGGCGGAGCCTATGGAAAAACGCCAGCAACGCGGCCTTTTTACGGTTCCTGGCCTTTTGCTGGCCTTTTGCTCACATGTTCTTTCCTGCGTTATCCCCTGATTCTGTGGATAACCGTATTACCGCCTTTGAGTGAGCTGATACCGCTCGCCGCAGCCGAACGACCGAGCGCAGCGAGTCAGTGAGCGAGGAAGCGGAAGAGCGCCCAATACGCAAACCGCCTCTCCCCGCGCGTTGGCCGATTCATTAATGCAGCTGGCACGACAGGTTTCCCGACTGGAAAGCGGGCAGTGAGCGCAACGCAATTAATGTGAGTTAGCTCACTCATTAGGCACCCCAGGCTTTACACTTTATGCTTCCGGCTCGTATGTTGTGTGGAATTGTGAGCGGATAACAATTTCACACAGGAAACAGCTATGACATGATTACGAATTAATTCGAGCTCGCCC

**Escape mutant: pRK5 myc-hDISC1 Escape mutant (escape mutation in DNA sequence is bold and highlighted in yellow)**

1822-ACC GAG GAG AT**C** AGA **AGC** **C**T**G** ACA- the sequence mutations introduced (indicated by bold and yellow) highlight resulted in silent mutations, which prevents the shRNA from binding but the amino acid sequence is intact.

GACATTGATTATTGACTAGTTATTAATAGTAATCAATTACGGGGTCATTAGTTCATAGCCCATATATGGAGTTCCGCGTTACATAACTTACGGTAAATGGCCCGCCTGGCTGACCGCCCAACGACCCCCGCCCATTGACGTCAATAATGACGTATGTTCCCATAGTAACGCCAATAGGGACTTTCCATTGACGTCAATGGGTGGAGTATTTACGGTAAACTGCCCACTTGGCAGTACATCAAGTGTATCATATGCCAAGTACGCCCCCTATTGACGTCAATGACGGTAAATGGCCCGCCTGGCATTATGCCCAGTACATGACCTTATGGGACTTTCCTACTTGGCAGTACATCTACGTATTAGTCATCGCTATTACCATGGTGATGCGGTTTTGGCAGTACATCAATGGGCGTGGATAGCGGTTTGACTCACGGGGATTTCCAAGTCTCCACCCCATTGACGTCAATGGGAGTTTGTTTTGGCACCAAAATCAACGGGACTTTCCAAAATGTCGTAACAACTCCGCCCCATTGACGCAAATGGGCGGTAGGCGTGTACGGTGGGAGGTCTATATAAGCAGAGCTCGTTTAGTGAACCGTCAGATCGCCTGGAGACGCCATCCACGCTGTTTTGACCTCCATAGAAGACACCGGGACCGATCCAGCCTCCGCGGCCGGGAACGGTGCATTGGAACGCGGATTCCCCGTGCCAAGAGTGACGTAAGTACCGCCTATAGAGTCTATAGGCCCACCCCCTTGGCTTCGTTAGAACGCGGCTACAATTAATACATAACCTTATGTATCATACACATACGATTTAGGTGACACTATAGAATAACATCCACTTTGCCTTTCTCTCCACAGGTGTCCACTCCCAGGTCCAACTGCACCTCGGTTCTATCGATTGAATTCCCGAACCGACAGTCGGTCTCTTCACCAAGGCCATTCGCGCCACCATGGAGCAAAAGCTCATTTCTGAGGAAGATCTCAATGGTGGTGGTGGTGGGTCGACCATGCCAGGCGGGGGTCCTCAGGGCGCCCCAGCCGCCGCCGGCGGCGGCGGCGTGAGCCACCGCGCAGGCAGCCGGGATTGCTTACCACCTGCAGCGTGCTTTCGGAGGCGGCGGCTGGCACGGAGGCCGGGCTACATGAGAAGCTCGACAGGGCCTGGGATCGGGTTCCTTTCCCCAGCAGTGGGCACACTGTTCCGGTTCCCAGGAGGGGTGTCTGGCGAGGAGTCCCACCACTCGGAGTCCAGGGCCAGACAGTGTGGCCTTGACTCGAGAGGCCTCTTGGTCCGGAGCCCTGTTTCCAAGAGTGCAGCAGCCCCTACTGTGACCTCTGTGAGAGGAACCTCGGCGCACTTTGGGATTCAGCTCAGAGGTGGCACCAGATTGCCTGACAGGCTTAGCTGGCCGTGTGGCCCTGGGAGTGCTGGGTGGCAGCAAGAGTTTGCAGCCATGGATAGTTCTGAGACCCTGGACGCCAGCTGGGAGGCAGCCTGCAGCGATGGAGCAAGGCGTGTCCGGGCAGCAGGCTCTCTGCCATCAGCAGAGTTGAGTAGCAACAGCTGCAGCCCTGGCTGTGGCCCTGAGGTCCCCCCAACCCCTCCTGGCTCTCACAGTGCCTTTACCTCAAGCTTTAGCTTTATTCGGCTCTCGCTTGGCTCTGCCGGGGAACGTGGAGAAGCAGAAGGCTGCCCACCATCCAGAGAGGCTGAGTCCCATTGCCAGAGCCCCCAGGAGATGGGAGCCAAAGCTGCCAGCTTGGACGGGCCTCACGAGGACCCGCGATGTCTCTCTCGGCCCTTCAGTCTCTTGGCTACACGGGTCTCTGCAGACTTGGCCCAGGCCGCAAGGAACAGCTCCAGGCCAGAGCGTGACATGCATTCTTTACCAGACATGGACCCTGGCTCCTCCAGTTCTCTGGATCCCTCACTGGCTGGCTGTGGTGGTGATGGGAGCAGCGGCTCAGGGGATGCCCACTCTTGGGACACCCTGCTCAGGAAATGGGAGCCAGTGCTGCGGGACTGCCTGCTGAGAAACCGGAGGCAGATGGAGGTAATATCCTTAAGATTAAAACTTCAGAAACTTCAGGAAGATGCAGTTGAGAATGATGATTATGATAAAGCTGAGACGTTACAACAAAGATTAGAAGACCTGGAACAAGAGAAAATCAGCCTGCACTTTCAACTTCCTTCAAGGCAGCCAGCTCTTAGCAGTTTCCTGGGTCACCTGGCAGCACAAGTCCAGGCTGCCTTGCGCCGTGGGGCCACTCAGCAGGCCAGCGGAGATGACACCCACACCCCACTGAGAATGGAGCCGAGGCTGTTGGAACCCACTGCTCAGGACAGCTTGCACGTGTCCATCACGAGACGAGACTGGCTTCTTCAGGAAAAGCAGCAGCTACAGAAAGAAATCGAAGCTCTCCAAGCAAGGATGTTTGTGCTGGAAGCCAAAGATCAACAGCTGAGAAGGGAAATAGAGGAGCAAGAGCAGCAACTCCAGTGGCAGGGCTGCGACCTGACCCCACTGGTGGGCCAGCTGTCCCTGGGTCAGCTGCAGGAGGTCAGCAAGGCCTTGCAGGACACCCTGGCCTCAGCCGGTCAGATTCCCTTCCATGCAGAGCCACCGGAAACCATAAGGAGCCTCCAGGAAAGAATAAAATCCCTCAACTTGTCACTTAAAGAAATCACTACTAAGGTGTGTATGAGTGAGAAATTCTGCAGCACCCTGAGGAAGAAAGTTAACGATATTGAAACCCAACTACCAGCCTTGCTTGAAGCCAAAATGCATGCCATATCAGGAAACCATTTCTGGACGGCTAAAGACCTCACCGAGGAGAT**C**AGA**AGCC**T**G**ACATCAGAGAGAGAAGGGCTGGAGGGACTCCTCAGCAAGCTGTTGGTGTTGAGTTCCAGGAATGTCAAAAAGCTGGGAAGTGTTAAAGAAGATTACAACAGACTGAGAAGAGAAGTGGAGCACCAGGAGACTGCCTATGAAACAAGTGTGAAGGAAAATACTATGAAGTACATGGAAACACTTAAGAATAAACTGTGCAGCTGCAAGTGTCCACTGCTTGGGAAAGTGTGGGAAGCTGACTTGGAAGCTTGTCGATTGCTTATCCAGAGCCTACAGCTCCAGGAAGCCAGGGGAAGCCTGTCTGTAGAAGATGAGAGGCAGATGGATGACTTAGAGGGAGCTGCTCCTCCTATTCCCCCCAGGCTCCACTCCGAGGATAAAAGGAAGACCCCTTTGAAGGTATTGGAAGAATGGAAGACTCACCTCATCCCCTCTCTGCACTGTGCTGGAGGTGAACAGAAAGAGGAATCTTACATCCTTTCTGCAGAACTTGGAGAAAAGTGTGAAGACATAGGCAAGAAGCTATTGTACTTGGAAGATCAACTTCACACAGCAATCCACAGTCATGATGAAGATCTCATTCAGTCTCTCAGGAGGGAGCTCCAGATGGTGAAGGAAACTCTGCAGGCCATGATCCTGCAGCTCCAGCCAGCAAAGGAGGCGGGAGAAAGAGAAGCTGCAGCTTCCTGCATGACAGCTGGTGTCCACGAAGCACAAGCCTGAGCGGCCGCTAAGTAAGTAAGGATCCCCAGCTTGGCCGCCATGGCCCAACTTGTTTATTGCAGCTTATAATGGTTACAAATAAAGCAATAGCATCACAAATTTCACAAATAAAGCATTTTTTTCACTGCATTCTAGTTGTGGTTTGTCCAAACTCATCAATGTATCTTATCATGTCTGGATCGGGAATTAATTCGGCGCAGCACCATGGCCTGAAATAACCTCTGAAAGAGGAACTTGGTTAGGTACCTTCTGAGGCGGAAAGAACCAGCTGTGGAATGTGTGTCAGTTAGGGTGTGGAAAGTCCCCAGGCTCCCCAGCAGGCAGAAGTATGCAAAGCATGCATCTCAATTAGTCAGCAACCAGGTGTGGAAAGTCCCCAGGCTCCCCAGCAGGCAGAAGTATGCAAAGCATGCATCTCAATTAGTCAGCAACCATAGTCCCGCCCCTAACTCCGCCCATCCCGCCCCTAACTCCGCCCAGTTCCGCCCATTCTCCGCCCCATGGCTGACTAATTTTTTTTATTTATGCAGAGGCCGAGGCCGCCTCGGCCTCTGAGCTATTCCAGAAGTAGTGAGGAGGCTTTTTTGGAGGCCTAGGCTTTTGCAAAAAGCTGTTAACAGCTTGGCACTGGCCGTCGTTTTACAACGTCGTGACTGGGAAAACCCTGGCGTTACCCAACTTAATCGCCTTGCAGCACATCCCCCTTTCGCCAGCTGGCGTAATAGCGAAGAGGCCCGCACCGATCGCCCTTCCCAACAGTTGCGCAGCCTGAATGGCGAATGGCGCCTGATGCGGTATTTTCTCCTTACGCATCTGTGCGGTATTTCACACCGCATACGTCAAAGCAACCATAGTACGCGCCCTGTAGCGGCGCATTAAGCGCGGCGGGTGTGGTGGTTACGCGCAGCGTGACCGCTACACTTGCCAGCGCCCTAGCGCCCGCTCCTTTCGCTTTCTTCCCTTCCTTTCTCGCCACGTTCGCCGGCTTTCCCCGTCAAGCTCTAAATCGGGGGCTCCCTTTAGGGTTCCGATTTAGTGCTTTACGGCACCTCGACCCCAAAAAACTTGATTTGGGTGATGGTTCACGTAGTGGGCCATCGCCCTGATAGACGGTTTTTCGCCCTTTGACGTTGGAGTCCACGTTCTTTAATAGTGGACTCTTGTTCCAAACTGGAACAACACTCAACCCTATCTCGGGCTATTCTTTTGATTTATAAGGGATTTTGCCGATTTCGGCCTATTGGTTAAAAAATGAGCTGATTTAACAAAAATTTAACGCGAATTTTAACAAAATATTAACGTTTACAATTTTATGGTGCACTCTCAGTACAATCTGCTCTGATGCCGCATAGTTAAGCCAGCCCCGACACCCGCCAACACCCGCTGACGCGCCCTGACGGGCTTGTCTGCTCCCGGCATCCGCTTACAGACAAGCTGTGACCGTCTCCGGGAGCTGCATGTGTCAGAGGTTTTCACCGTCATCACCGAAACGCGCGAGACGAAAGGGCCTCGTGATACGCCTATTTTTATAGGTTAATGTCATGATAATAATGGTTTCTTAGACGTCAGGTGGCACTTTTCGGGGAAATGTGCGCGGAACCCCTATTTGTTTATTTTTCTAAATACATTCAAATATGTATCCGCTCATGAGACAATAACCCTGATAAATGCTTCAATAATATTGAAAAAGGAAGAGTATGAGTATTCAACATTTCCGTGTCGCCCTTATTCCCTTTTTTGCGGCATTTTGCCTTCCTGTTTTTGCTCACCCAGAAACGCTGGTGAAAGTAAAAGATGCTGAAGATCAGTTGGGTGCACGAGTGGGTTACATCGAACTGGATCTCAACAGCGGTAAGATCCTTGAGAGTTTTCGCCCCGAAGAACGTTTTCCAATGATGAGCACTTTTAAAGTTCTGCTATGTGGCGCGGTATTATCCCGTATTGACGCCGGGCAAGAGCAACTCGGTCGCCGCATACACTATTCTCAGAATGACTTGGTTGAGTACTCACCAGTCACAGAAAAGCATCTTACGGATGGCATGACAGTAAGAGAATTATGCAGTGCTGCCATAACCATGAGTGATAACACTGCGGCCAACTTACTTCTGACAACGATCGGAGGACCGAAGGAGCTAACCGCTTTTTTGCACAACATGGGGGATCATGTAACTCGCCTTGATCGTTGGGAACCGGAGCTGAATGAAGCCATACCAAACGACGAGCGTGACACCACGATGCCTGTAGCAATGGCAACAACGTTGCGCAAACTATTAACTGGCGAACTACTTACTCTAGCTTCCCGGCAACAATTAATAGACTGGATGGAGGCGGATAAAGTTGCAGGACCACTTCTGCGCTCGGCCCTTCCGGCTGGCTGGTTTATTGCTGATAAATCTGGAGCCGGTGAGCGTGGGTCTCGCGGTATCATTGCAGCACTGGGGCCAGATGGTAAGCCCTCCCGTATCGTAGTTATCTACACGACGGGGAGTCAGGCAACTATGGATGAACGAAATAGACAGATCGCTGAGATAGGTGCCTCACTGATTAAGCATTGGTAACTGTCAGACCAAGTTTACTCATATATACTTTAGATTGATTTAAAACTTCATTTTTAATTTAAAAGGATCTAGGTGAAGATCCTTTTTGATAATCTCATGACCAAAATCCCTTAACGTGAGTTTTCGTTCCACTGAGCGTCAGACCCCGTAGAAAAGATCAAAGGATCTTCTTGAGATCCTTTTTTTCTGCGCGTAATCTGCTGCTTGCAAACAAAAAAACCACCGCTACCAGCGGTGGTTTGTTTGCCGGATCAAGAGCTACCAACTCTTTTTCCGAAGGTAACTGGCTTCAGCAGAGCGCAGATACCAAATACTGTTCTTCTAGTGTAGCCGTAGTTAGGCCACCACTTCAAGAACTCTGTAGCACCGCCTACATACCTCGCTCTGCTAATCCTGTTACCAGTGGCTGCTGCCAGTGGCGATAAGTCGTGTCTTACCGGGTTGGACTCAAGACGATAGTTACCGGATAAGGCGCAGCGGTCGGGCTGAACGGGGGGTTCGTGCACACAGCCCAGCTTGGAGCGAACGACCTACACCGAACTGAGATACCTACAGCGTGAGCTATGAGAAAGCGCCACGCTTCCCGAAGGGAGAAAGGCGGACAGGTATCCGGTAAGCGGCAGGGTCGGAACAGGAGAGCGCACGAGGGAGCTTCCAGGGGGAAACGCCTGGTATCTTTATAGTCCTGTCGGGTTTCGCCACCTCTGACTTGAGCGTCGATTTTTGTGATGCTCGTCAGGGGGGCGGAGCCTATGGAAAAACGCCAGCAACGCGGCCTTTTTACGGTTCCTGGCCTTTTGCTGGCCTTTTGCTCACATGTTCTTTCCTGCGTTATCCCCTGATTCTGTGGATAACCGTATTACCGCCTTTGAGTGAGCTGATACCGCTCGCCGCAGCCGAACGACCGAGCGCAGCGAGTCAGTGAGCGAGGAAGCGGAAGAGCGCCCAATACGCAAACCGCCTCTCCCCGCGCGTTGGCCGATTCATTAATGCAGCTGGCACGACAGGTTTCCCGACTGGAAAGCGGGCAGTGAGCGCAACGCAATTAATGTGAGTTAGCTCACTCATTAGGCACCCCAGGCTTTACACTTTATGCTTCCGGCTCGTATGTTGTGTGGAATTGTGAGCGGATAACAATTTCACACAGGAAACAGCTATGACATGATTACGAATTAATTCGAGCTCGCCC

**F to A mutant: pRK5 myc-hDISC1 F206A, F210A, F212A**

**F to A mutations shown in bold and red highlighted**

GACATTGATTATTGACTAGTTATTAATAGTAATCAATTACGGGGTCATTAGTTCATAGCCCATATATGGAGTTCCGCGTTACATAACTTACGGTAAATGGCCCGCCTGGCTGACCGCCCAACGACCCCCGCCCATTGACGTCAATAATGACGTATGTTCCCATAGTAACGCCAATAGGGACTTTCCATTGACGTCAATGGGTGGAGTATTTACGGTAAACTGCCCACTTGGCAGTACATCAAGTGTATCATATGCCAAGTACGCCCCCTATTGACGTCAATGACGGTAAATGGCCCGCCTGGCATTATGCCCAGTACATGACCTTATGGGACTTTCCTACTTGGCAGTACATCTACGTATTAGTCATCGCTATTACCATGGTGATGCGGTTTTGGCAGTACATCAATGGGCGTGGATAGCGGTTTGACTCACGGGGATTTCCAAGTCTCCACCCCATTGACGTCAATGGGAGTTTGTTTTGGCACCAAAATCAACGGGACTTTCCAAAATGTCGTAACAACTCCGCCCCATTGACGCAAATGGGCGGTAGGCGTGTACGGTGGGAGGTCTATATAAGCAGAGCTCGTTTAGTGAACCGTCAGATCGCCTGGAGACGCCATCCACGCTGTTTTGACCTCCATAGAAGACACCGGGACCGATCCAGCCTCCGCGGCCGGGAACGGTGCATTGGAACGCGGATTCCCCGTGCCAAGAGTGACGTAAGTACCGCCTATAGAGTCTATAGGCCCACCCCCTTGGCTTCGTTAGAACGCGGCTACAATTAATACATAACCTTATGTATCATACACATACGATTTAGGTGACACTATAGAATAACATCCACTTTGCCTTTCTCTCCACAGGTGTCCACTCCCAGGTCCAACTGCACCTCGGTTCTATCGATTGAATTCCCGAACCGACAGTCGGTCTCTTCACCAAGGCCATTCGCGCCACCATGGAGCAAAAGCTCATTTCTGAGGAAGATCTCAATGGTGGTGGTGGTGGGTCGACCATGCCAGGCGGGGGTCCTCAGGGCGCCCCAGCCGCCGCCGGCGGCGGCGGCGTGAGCCACCGCGCAGGCAGCCGGGATTGCTTACCACCTGCAGCGTGCTTTCGGAGGCGGCGGCTGGCACGGAGGCCGGGCTACATGAGAAGCTCGACAGGGCCTGGGATCGGGTTCCTTTCCCCAGCAGTGGGCACACTGTTCCGGTTCCCAGGAGGGGTGTCTGGCGAGGAGTCCCACCACTCGGAGTCCAGGGCCAGACAGTGTGGCCTTGACTCGAGAGGCCTCTTGGTCCGGAGCCCTGTTTCCAAGAGTGCAGCAGCCCCTACTGTGACCTCTGTGAGAGGAACCTCGGCGCACTTTGGGATTCAGCTCAGAGGTGGCACCAGATTGCCTGACAGGCTTAGCTGGCCGTGTGGCCCTGGGAGTGCTGGGTGGCAGCAAGAGTTTGCAGCCATGGATAGTTCTGAGACCCTGGACGCCAGCTGGGAGGCAGCCTGCAGCGATGGAGCAAGGCGTGTCCGGGCAGCAGGCTCTCTGCCATCAGCAGAGTTGAGTAGCAACAGCTGCAGCCCTGGCTGTGGCCCTGAGGTCCCCCCAACCCCTCCTGGCTCTCACAGTGCC**GCA**ACCAGCTCC**GCC**TCC**GCC**ATTCGGCTCTCGCTTGGCTCTGCCGGGGAACGTGGAGAAGCAGAAGGCTGCCCACCATCCAGAGAGGCTGAGTCCCATTGCCAGAGCCCCCAGGAGATGGGAGCCAAAGCTGCCAGCTTGGACGGGCCTCACGAGGACCCGCGATGTCTCTCTCGGCCCTTCAGTCTCTTGGCTACACGGGTCTCTGCAGACTTGGCCCAGGCCGCAAGGAACAGCTCCAGGCCAGAGCGTGACATGCATTCTTTACCAGACATGGACCCTGGCTCCTCCAGTTCTCTGGATCCCTCACTGGCTGGCTGTGGTGGTGATGGGAGCAGCGGCTCAGGGGATGCCCACTCTTGGGACACCCTGCTCAGGAAATGGGAGCCAGTGCTGCGGGACTGCCTGCTGAGAAACCGGAGGCAGATGGAGGTAATATCCTTAAGATTAAAACTTCAGAAACTTCAGGAAGATGCAGTTGAGAATGATGATTATGATAAAGCTGAGACGTTACAACAAAGATTAGAAGACCTGGAACAAGAGAAAATCAGCCTGCACTTTCAACTTCCTTCAAGGCAGCCAGCTCTTAGCAGTTTCCTGGGTCACCTGGCAGCACAAGTCCAGGCTGCCTTGCGCCGTGGGGCCACTCAGCAGGCCAGCGGAGATGACACCCACACCCCACTGAGAATGGAGCCGAGGCTGTTGGAACCCACTGCTCAGGACAGCTTGCACGTGTCCATCACGAGACGAGACTGGCTTCTTCAGGAAAAGCAGCAGCTACAGAAAGAAATCGAAGCTCTCCAAGCAAGGATGTTTGTGCTGGAAGCCAAAGATCAACAGCTGAGAAGGGAAATAGAGGAGCAAGAGCAGCAACTCCAGTGGCAGGGCTGCGACCTGACCCCACTGGTGGGCCAGCTGTCCCTGGGTCAGCTGCAGGAGGTCAGCAAGGCCTTGCAGGACACCCTGGCCTCAGCCGGTCAGATTCCCTTCCATGCAGAGCCACCGGAAACCATAAGGAGCCTCCAGGAAAGAATAAAATCCCTCAACTTGTCACTTAAAGAAATCACTACTAAGGTGTGTATGAGTGAGAAATTCTGCAGCACCCTGAGGAAGAAAGTTAACGATATTGAAACCCAACTACCAGCCTTGCTTGAAGCCAAAATGCATGCCATATCAGGAAACCATTTCTGGACGGCTAAAGACCTCACCGAGGAGATTAGATCATTAACATCAGAGAGAGAAGGGCTGGAGGGACTCCTCAGCAAGCTGTTGGTGTTGAGTTCCAGGAATGTCAAAAAGCTGGGAAGTGTTAAAGAAGATTACAACAGACTGAGAAGAGAAGTGGAGCACCAGGAGACTGCCTATGAAACAAGTGTGAAGGAAAATACTATGAAGTACATGGAAACACTTAAGAATAAACTGTGCAGCTGCAAGTGTCCACTGCTTGGGAAAGTGTGGGAAGCTGACTTGGAAGCTTGTCGATTGCTTATCCAGAGCCTACAGCTCCAGGAAGCCAGGGGAAGCCTGTCTGTAGAAGATGAGAGGCAGATGGATGACTTAGAGGGAGCTGCTCCTCCTATTCCCCCCAGGCTCCACTCCGAGGATAAAAGGAAGACCCCTTTGAAGGTATTGGAAGAATGGAAGACTCACCTCATCCCCTCTCTGCACTGTGCTGGAGGTGAACAGAAAGAGGAATCTTACATCCTTTCTGCAGAACTTGGAGAAAAGTGTGAAGACATAGGCAAGAAGCTATTGTACTTGGAAGATCAACTTCACACAGCAATCCACAGTCATGATGAAGATCTCATTCAGTCTCTCAGGAGGGAGCTCCAGATGGTGAAGGAAACTCTGCAGGCCATGATCCTGCAGCTCCAGCCAGCAAAGGAGGCGGGAGAAAGAGAAGCTGCAGCTTCCTGCATGACAGCTGGTGTCCACGAAGCACAAGCCTGAGCGGCCGCTAAGTAAGTAAGGATCCCCAGCTTGGCCGCCATGGCCCAACTTGTTTATTGCAGCTTATAATGGTTACAAATAAAGCAATAGCATCACAAATTTCACAAATAAAGCATTTTTTTCACTGCATTCTAGTTGTGGTTTGTCCAAACTCATCAATGTATCTTATCATGTCTGGATCGGGAATTAATTCGGCGCAGCACCATGGCCTGAAATAACCTCTGAAAGAGGAACTTGGTTAGGTACCTTCTGAGGCGGAAAGAACCAGCTGTGGAATGTGTGTCAGTTAGGGTGTGGAAAGTCCCCAGGCTCCCCAGCAGGCAGAAGTATGCAAAGCATGCATCTCAATTAGTCAGCAACCAGGTGTGGAAAGTCCCCAGGCTCCCCAGCAGGCAGAAGTATGCAAAGCATGCATCTCAATTAGTCAGCAACCATAGTCCCGCCCCTAACTCCGCCCATCCCGCCCCTAACTCCGCCCAGTTCCGCCCATTCTCCGCCCCATGGCTGACTAATTTTTTTTATTTATGCAGAGGCCGAGGCCGCCTCGGCCTCTGAGCTATTCCAGAAGTAGTGAGGAGGCTTTTTTGGAGGCCTAGGCTTTTGCAAAAAGCTGTTAACAGCTTGGCACTGGCCGTCGTTTTACAACGTCGTGACTGGGAAAACCCTGGCGTTACCCAACTTAATCGCCTTGCAGCACATCCCCCTTTCGCCAGCTGGCGTAATAGCGAAGAGGCCCGCACCGATCGCCCTTCCCAACAGTTGCGCAGCCTGAATGGCGAATGGCGCCTGATGCGGTATTTTCTCCTTACGCATCTGTGCGGTATTTCACACCGCATACGTCAAAGCAACCATAGTACGCGCCCTGTAGCGGCGCATTAAGCGCGGCGGGTGTGGTGGTTACGCGCAGCGTGACCGCTACACTTGCCAGCGCCCTAGCGCCCGCTCCTTTCGCTTTCTTCCCTTCCTTTCTCGCCACGTTCGCCGGCTTTCCCCGTCAAGCTCTAAATCGGGGGCTCCCTTTAGGGTTCCGATTTAGTGCTTTACGGCACCTCGACCCCAAAAAACTTGATTTGGGTGATGGTTCACGTAGTGGGCCATCGCCCTGATAGACGGTTTTTCGCCCTTTGACGTTGGAGTCCACGTTCTTTAATAGTGGACTCTTGTTCCAAACTGGAACAACACTCAACCCTATCTCGGGCTATTCTTTTGATTTATAAGGGATTTTGCCGATTTCGGCCTATTGGTTAAAAAATGAGCTGATTTAACAAAAATTTAACGCGAATTTTAACAAAATATTAACGTTTACAATTTTATGGTGCACTCTCAGTACAATCTGCTCTGATGCCGCATAGTTAAGCCAGCCCCGACACCCGCCAACACCCGCTGACGCGCCCTGACGGGCTTGTCTGCTCCCGGCATCCGCTTACAGACAAGCTGTGACCGTCTCCGGGAGCTGCATGTGTCAGAGGTTTTCACCGTCATCACCGAAACGCGCGAGACGAAAGGGCCTCGTGATACGCCTATTTTTATAGGTTAATGTCATGATAATAATGGTTTCTTAGACGTCAGGTGGCACTTTTCGGGGAAATGTGCGCGGAACCCCTATTTGTTTATTTTTCTAAATACATTCAAATATGTATCCGCTCATGAGACAATAACCCTGATAAATGCTTCAATAATATTGAAAAAGGAAGAGTATGAGTATTCAACATTTCCGTGTCGCCCTTATTCCCTTTTTTGCGGCATTTTGCCTTCCTGTTTTTGCTCACCCAGAAACGCTGGTGAAAGTAAAAGATGCTGAAGATCAGTTGGGTGCACGAGTGGGTTACATCGAACTGGATCTCAACAGCGGTAAGATCCTTGAGAGTTTTCGCCCCGAAGAACGTTTTCCAATGATGAGCACTTTTAAAGTTCTGCTATGTGGCGCGGTATTATCCCGTATTGACGCCGGGCAAGAGCAACTCGGTCGCCGCATACACTATTCTCAGAATGACTTGGTTGAGTACTCACCAGTCACAGAAAAGCATCTTACGGATGGCATGACAGTAAGAGAATTATGCAGTGCTGCCATAACCATGAGTGATAACACTGCGGCCAACTTACTTCTGACAACGATCGGAGGACCGAAGGAGCTAACCGCTTTTTTGCACAACATGGGGGATCATGTAACTCGCCTTGATCGTTGGGAACCGGAGCTGAATGAAGCCATACCAAACGACGAGCGTGACACCACGATGCCTGTAGCAATGGCAACAACGTTGCGCAAACTATTAACTGGCGAACTACTTACTCTAGCTTCCCGGCAACAATTAATAGACTGGATGGAGGCGGATAAAGTTGCAGGACCACTTCTGCGCTCGGCCCTTCCGGCTGGCTGGTTTATTGCTGATAAATCTGGAGCCGGTGAGCGTGGGTCTCGCGGTATCATTGCAGCACTGGGGCCAGATGGTAAGCCCTCCCGTATCGTAGTTATCTACACGACGGGGAGTCAGGCAACTATGGATGAACGAAATAGACAGATCGCTGAGATAGGTGCCTCACTGATTAAGCATTGGTAACTGTCAGACCAAGTTTACTCATATATACTTTAGATTGATTTAAAACTTCATTTTTAATTTAAAAGGATCTAGGTGAAGATCCTTTTTGATAATCTCATGACCAAAATCCCTTAACGTGAGTTTTCGTTCCACTGAGCGTCAGACCCCGTAGAAAAGATCAAAGGATCTTCTTGAGATCCTTTTTTTCTGCGCGTAATCTGCTGCTTGCAAACAAAAAAACCACCGCTACCAGCGGTGGTTTGTTTGCCGGATCAAGAGCTACCAACTCTTTTTCCGAAGGTAACTGGCTTCAGCAGAGCGCAGATACCAAATACTGTTCTTCTAGTGTAGCCGTAGTTAGGCCACCACTTCAAGAACTCTGTAGCACCGCCTACATACCTCGCTCTGCTAATCCTGTTACCAGTGGCTGCTGCCAGTGGCGATAAGTCGTGTCTTACCGGGTTGGACTCAAGACGATAGTTACCGGATAAGGCGCAGCGGTCGGGCTGAACGGGGGGTTCGTGCACACAGCCCAGCTTGGAGCGAACGACCTACACCGAACTGAGATACCTACAGCGTGAGCTATGAGAAAGCGCCACGCTTCCCGAAGGGAGAAAGGCGGACAGGTATCCGGTAAGCGGCAGGGTCGGAACAGGAGAGCGCACGAGGGAGCTTCCAGGGGGAAACGCCTGGTATCTTTATAGTCCTGTCGGGTTTCGCCACCTCTGACTTGAGCGTCGATTTTTGTGATGCTCGTCAGGGGGGCGGAGCCTATGGAAAAACGCCAGCAACGCGGCCTTTTTACGGTTCCTGGCCTTTTGCTGGCCTTTTGCTCACATGTTCTTTCCTGCGTTATCCCCTGATTCTGTGGATAACCGTATTACCGCCTTTGAGTGAGCTGATACCGCTCGCCGCAGCCGAACGACCGAGCGCAGCGAGTCAGTGAGCGAGGAAGCGGAAGAGCGCCCAATACGCAAACCGCCTCTCCCCGCGCGTTGGCCGATTCATTAATGCAGCTGGCACGACAGGTTTCCCGACTGGAAAGCGGGCAGTGAGCGCAACGCAATTAATGTGAGTTAGCTCACTCATTAGGCACCCCAGGCTTTACACTTTATGCTTCCGGCTCGTATGTTGTGTGGAATTGTGAGCGGATAACAATTTCACACAGGAAACAGCTATGACATGATTACGAATTAATTCGAGCTCGCCC

**F to A Escape mutant: pRK5 myc-DISC1 Escape mutant F206A, F210A, F212A**

**(escape mutation in DNA sequence is bold and highlighted in yellow, F to A mutations shown in bold and red highlighted)**

GACATTGATTATTGACTAGTTATTAATAGTAATCAATTACGGGGTCATTAGTTCATAGCCCATATATGGAGTTCCGCGTTACATAACTTACGGTAAATGGCCCGCCTGGCTGACCGCCCAACGACCCCCGCCCATTGACGTCAATAATGACGTATGTTCCCATAGTAACGCCAATAGGGACTTTCCATTGACGTCAATGGGTGGAGTATTTACGGTAAACTGCCCACTTGGCAGTACATCAAGTGTATCATATGCCAAGTACGCCCCCTATTGACGTCAATGACGGTAAATGGCCCGCCTGGCATTATGCCCAGTACATGACCTTATGGGACTTTCCTACTTGGCAGTACATCTACGTATTAGTCATCGCTATTACCATGGTGATGCGGTTTTGGCAGTACATCAATGGGCGTGGATAGCGGTTTGACTCACGGGGATTTCCAAGTCTCCACCCCATTGACGTCAATGGGAGTTTGTTTTGGCACCAAAATCAACGGGACTTTCCAAAATGTCGTAACAACTCCGCCCCATTGACGCAAATGGGCGGTAGGCGTGTACGGTGGGAGGTCTATATAAGCAGAGCTCGTTTAGTGAACCGTCAGATCGCCTGGAGACGCCATCCACGCTGTTTTGACCTCCATAGAAGACACCGGGACCGATCCAGCCTCCGCGGCCGGGAACGGTGCATTGGAACGCGGATTCCCCGTGCCAAGAGTGACGTAAGTACCGCCTATAGAGTCTATAGGCCCACCCCCTTGGCTTCGTTAGAACGCGGCTACAATTAATACATAACCTTATGTATCATACACATACGATTTAGGTGACACTATAGAATAACATCCACTTTGCCTTTCTCTCCACAGGTGTCCACTCCCAGGTCCAACTGCACCTCGGTTCTATCGATTGAATTCCCGAACCGACAGTCGGTCTCTTCACCAAGGCCATTCGCGCCACCATGGAGCAAAAGCTCATTTCTGAGGAAGATCTCAATGGTGGTGGTGGTGGGTCGACCATGCCAGGCGGGGGTCCTCAGGGCGCCCCAGCCGCCGCCGGCGGCGGCGGCGTGAGCCACCGCGCAGGCAGCCGGGATTGCTTACCACCTGCAGCGTGCTTTCGGAGGCGGCGGCTGGCACGGAGGCCGGGCTACATGAGAAGCTCGACAGGGCCTGGGATCGGGTTCCTTTCCCCAGCAGTGGGCACACTGTTCCGGTTCCCAGGAGGGGTGTCTGGCGAGGAGTCCCACCACTCGGAGTCCAGGGCCAGACAGTGTGGCCTTGACTCGAGAGGCCTCTTGGTCCGGAGCCCTGTTTCCAAGAGTGCAGCAGCCCCTACTGTGACCTCTGTGAGAGGAACCTCGGCGCACTTTGGGATTCAGCTCAGAGGTGGCACCAGATTGCCTGACAGGCTTAGCTGGCCGTGTGGCCCTGGGAGTGCTGGGTGGCAGCAAGAGTTTGCAGCCATGGATAGTTCTGAGACCCTGGACGCCAGCTGGGAGGCAGCCTGCAGCGATGGAGCAAGGCGTGTCCGGGCAGCAGGCTCTCTGCCATCAGCAGAGTTGAGTAGCAACAGCTGCAGCCCTGGCTGTGGCCCTGAGGTCCCCCCAACCCCTCCTGGCTCTCACAGTGCC**GCA**ACCAGCTCC**GCC**TCC**GCC**ATTCGGCTCTCGCTTGGCTCTGCCGGGGAACGTGGAGAAGCAGAAGGCTGCCCACCATCCAGAGAGGCTGAGTCCCATTGCCAGAGCCCCCAGGAGATGGGAGCCAAAGCTGCCAGCTTGGACGGGCCTCACGAGGACCCGCGATGTCTCTCTCGGCCCTTCAGTCTCTTGGCTACACGGGTCTCTGCAGACTTGGCCCAGGCCGCAAGGAACAGCTCCAGGCCAGAGCGTGACATGCATTCTTTACCAGACATGGACCCTGGCTCCTCCAGTTCTCTGGATCCCTCACTGGCTGGCTGTGGTGGTGATGGGAGCAGCGGCTCAGGGGATGCCCACTCTTGGGACACCCTGCTCAGGAAATGGGAGCCAGTGCTGCGGGACTGCCTGCTGAGAAACCGGAGGCAGATGGAGGTAATATCCTTAAGATTAAAACTTCAGAAACTTCAGGAAGATGCAGTTGAGAATGATGATTATGATAAAGCTGAGACGTTACAACAAAGATTAGAAGACCTGGAACAAGAGAAAATCAGCCTGCACTTTCAACTTCCTTCAAGGCAGCCAGCTCTTAGCAGTTTCCTGGGTCACCTGGCAGCACAAGTCCAGGCTGCCTTGCGCCGTGGGGCCACTCAGCAGGCCAGCGGAGATGACACCCACACCCCACTGAGAATGGAGCCGAGGCTGTTGGAACCCACTGCTCAGGACAGCTTGCACGTGTCCATCACGAGACGAGACTGGCTTCTTCAGGAAAAGCAGCAGCTACAGAAAGAAATCGAAGCTCTCCAAGCAAGGATGTTTGTGCTGGAAGCCAAAGATCAACAGCTGAGAAGGGAAATAGAGGAGCAAGAGCAGCAACTCCAGTGGCAGGGCTGCGACCTGACCCCACTGGTGGGCCAGCTGTCCCTGGGTCAGCTGCAGGAGGTCAGCAAGGCCTTGCAGGACACCCTGGCCTCAGCCGGTCAGATTCCCTTCCATGCAGAGCCACCGGAAACCATAAGGAGCCTCCAGGAAAGAATAAAATCCCTCAACTTGTCACTTAAAGAAATCACTACTAAGGTGTGTATGAGTGAGAAATTCTGCAGCACCCTGAGGAAGAAAGTTAACGATATTGAAACCCAACTACCAGCCTTGCTTGAAGCCAAAATGCATGCCATATCAGGAAACCATTTCTGGACGGCTAAAGACCTCACCGAGGAGAT**C**AGA**AGCC**T**G**ACATCAGAGAGAGAAGGGCTGGAGGGACTCCTCAGCAAGCTGTTGGTGTTGAGTTCCAGGAATGTCAAAAAGCTGGGAAGTGTTAAAGAAGATTACAACAGACTGAGAAGAGAAGTGGAGCACCAGGAGACTGCCTATGAAACAAGTGTGAAGGAAAATACTATGAAGTACATGGAAACACTTAAGAATAAACTGTGCAGCTGCAAGTGTCCACTGCTTGGGAAAGTGTGGGAAGCTGACTTGGAAGCTTGTCGATTGCTTATCCAGAGCCTACAGCTCCAGGAAGCCAGGGGAAGCCTGTCTGTAGAAGATGAGAGGCAGATGGATGACTTAGAGGGAGCTGCTCCTCCTATTCCCCCCAGGCTCCACTCCGAGGATAAAAGGAAGACCCCTTTGAAGGTATTGGAAGAATGGAAGACTCACCTCATCCCCTCTCTGCACTGTGCTGGAGGTGAACAGAAAGAGGAATCTTACATCCTTTCTGCAGAACTTGGAGAAAAGTGTGAAGACATAGGCAAGAAGCTATTGTACTTGGAAGATCAACTTCACACAGCAATCCACAGTCATGATGAAGATCTCATTCAGTCTCTCAGGAGGGAGCTCCAGATGGTGAAGGAAACTCTGCAGGCCATGATCCTGCAGCTCCAGCCAGCAAAGGAGGCGGGAGAAAGAGAAGCTGCAGCTTCCTGCATGACAGCTGGTGTCCACGAAGCACAAGCCTGAGCGGCCGCTAAGTAAGTAAGGATCCCCAGCTTGGCCGCCATGGCCCAACTTGTTTATTGCAGCTTATAATGGTTACAAATAAAGCAATAGCATCACAAATTTCACAAATAAAGCATTTTTTTCACTGCATTCTAGTTGTGGTTTGTCCAAACTCATCAATGTATCTTATCATGTCTGGATCGGGAATTAATTCGGCGCAGCACCATGGCCTGAAATAACCTCTGAAAGAGGAACTTGGTTAGGTACCTTCTGAGGCGGAAAGAACCAGCTGTGGAATGTGTGTCAGTTAGGGTGTGGAAAGTCCCCAGGCTCCCCAGCAGGCAGAAGTATGCAAAGCATGCATCTCAATTAGTCAGCAACCAGGTGTGGAAAGTCCCCAGGCTCCCCAGCAGGCAGAAGTATGCAAAGCATGCATCTCAATTAGTCAGCAACCATAGTCCCGCCCCTAACTCCGCCCATCCCGCCCCTAACTCCGCCCAGTTCCGCCCATTCTCCGCCCCATGGCTGACTAATTTTTTTTATTTATGCAGAGGCCGAGGCCGCCTCGGCCTCTGAGCTATTCCAGAAGTAGTGAGGAGGCTTTTTTGGAGGCCTAGGCTTTTGCAAAAAGCTGTTAACAGCTTGGCACTGGCCGTCGTTTTACAACGTCGTGACTGGGAAAACCCTGGCGTTACCCAACTTAATCGCCTTGCAGCACATCCCCCTTTCGCCAGCTGGCGTAATAGCGAAGAGGCCCGCACCGATCGCCCTTCCCAACAGTTGCGCAGCCTGAATGGCGAATGGCGCCTGATGCGGTATTTTCTCCTTACGCATCTGTGCGGTATTTCACACCGCATACGTCAAAGCAACCATAGTACGCGCCCTGTAGCGGCGCATTAAGCGCGGCGGGTGTGGTGGTTACGCGCAGCGTGACCGCTACACTTGCCAGCGCCCTAGCGCCCGCTCCTTTCGCTTTCTTCCCTTCCTTTCTCGCCACGTTCGCCGGCTTTCCCCGTCAAGCTCTAAATCGGGGGCTCCCTTTAGGGTTCCGATTTAGTGCTTTACGGCACCTCGACCCCAAAAAACTTGATTTGGGTGATGGTTCACGTAGTGGGCCATCGCCCTGATAGACGGTTTTTCGCCCTTTGACGTTGGAGTCCACGTTCTTTAATAGTGGACTCTTGTTCCAAACTGGAACAACACTCAACCCTATCTCGGGCTATTCTTTTGATTTATAAGGGATTTTGCCGATTTCGGCCTATTGGTTAAAAAATGAGCTGATTTAACAAAAATTTAACGCGAATTTTAACAAAATATTAACGTTTACAATTTTATGGTGCACTCTCAGTACAATCTGCTCTGATGCCGCATAGTTAAGCCAGCCCCGACACCCGCCAACACCCGCTGACGCGCCCTGACGGGCTTGTCTGCTCCCGGCATCCGCTTACAGACAAGCTGTGACCGTCTCCGGGAGCTGCATGTGTCAGAGGTTTTCACCGTCATCACCGAAACGCGCGAGACGAAAGGGCCTCGTGATACGCCTATTTTTATAGGTTAATGTCATGATAATAATGGTTTCTTAGACGTCAGGTGGCACTTTTCGGGGAAATGTGCGCGGAACCCCTATTTGTTTATTTTTCTAAATACATTCAAATATGTATCCGCTCATGAGACAATAACCCTGATAAATGCTTCAATAATATTGAAAAAGGAAGAGTATGAGTATTCAACATTTCCGTGTCGCCCTTATTCCCTTTTTTGCGGCATTTTGCCTTCCTGTTTTTGCTCACCCAGAAACGCTGGTGAAAGTAAAAGATGCTGAAGATCAGTTGGGTGCACGAGTGGGTTACATCGAACTGGATCTCAACAGCGGTAAGATCCTTGAGAGTTTTCGCCCCGAAGAACGTTTTCCAATGATGAGCACTTTTAAAGTTCTGCTATGTGGCGCGGTATTATCCCGTATTGACGCCGGGCAAGAGCAACTCGGTCGCCGCATACACTATTCTCAGAATGACTTGGTTGAGTACTCACCAGTCACAGAAAAGCATCTTACGGATGGCATGACAGTAAGAGAATTATGCAGTGCTGCCATAACCATGAGTGATAACACTGCGGCCAACTTACTTCTGACAACGATCGGAGGACCGAAGGAGCTAACCGCTTTTTTGCACAACATGGGGGATCATGTAACTCGCCTTGATCGTTGGGAACCGGAGCTGAATGAAGCCATACCAAACGACGAGCGTGACACCACGATGCCTGTAGCAATGGCAACAACGTTGCGCAAACTATTAACTGGCGAACTACTTACTCTAGCTTCCCGGCAACAATTAATAGACTGGATGGAGGCGGATAAAGTTGCAGGACCACTTCTGCGCTCGGCCCTTCCGGCTGGCTGGTTTATTGCTGATAAATCTGGAGCCGGTGAGCGTGGGTCTCGCGGTATCATTGCAGCACTGGGGCCAGATGGTAAGCCCTCCCGTATCGTAGTTATCTACACGACGGGGAGTCAGGCAACTATGGATGAACGAAATAGACAGATCGCTGAGATAGGTGCCTCACTGATTAAGCATTGGTAACTGTCAGACCAAGTTTACTCATATATACTTTAGATTGATTTAAAACTTCATTTTTAATTTAAAAGGATCTAGGTGAAGATCCTTTTTGATAATCTCATGACCAAAATCCCTTAACGTGAGTTTTCGTTCCACTGAGCGTCAGACCCCGTAGAAAAGATCAAAGGATCTTCTTGAGATCCTTTTTTTCTGCGCGTAATCTGCTGCTTGCAAACAAAAAAACCACCGCTACCAGCGGTGGTTTGTTTGCCGGATCAAGAGCTACCAACTCTTTTTCCGAAGGTAACTGGCTTCAGCAGAGCGCAGATACCAAATACTGTTCTTCTAGTGTAGCCGTAGTTAGGCCACCACTTCAAGAACTCTGTAGCACCGCCTACATACCTCGCTCTGCTAATCCTGTTACCAGTGGCTGCTGCCAGTGGCGATAAGTCGTGTCTTACCGGGTTGGACTCAAGACGATAGTTACCGGATAAGGCGCAGCGGTCGGGCTGAACGGGGGGTTCGTGCACACAGCCCAGCTTGGAGCGAACGACCTACACCGAACTGAGATACCTACAGCGTGAGCTATGAGAAAGCGCCACGCTTCCCGAAGGGAGAAAGGCGGACAGGTATCCGGTAAGCGGCAGGGTCGGAACAGGAGAGCGCACGAGGGAGCTTCCAGGGGGAAACGCCTGGTATCTTTATAGTCCTGTCGGGTTTCGCCACCTCTGACTTGAGCGTCGATTTTTGTGATGCTCGTCAGGGGGGCGGAGCCTATGGAAAAACGCCAGCAACGCGGCCTTTTTACGGTTCCTGGCCTTTTGCTGGCCTTTTGCTCACATGTTCTTTCCTGCGTTATCCCCTGATTCTGTGGATAACCGTATTACCGCCTTTGAGTGAGCTGATACCGCTCGCCGCAGCCGAACGACCGAGCGCAGCGAGTCAGTGAGCGAGGAAGCGGAAGAGCGCCCAATACGCAAACCGCCTCTCCCCGCGCGTTGGCCGATTCATTAATGCAGCTGGCACGACAGGTTTCCCGACTGGAAAGCGGGCAGTGAGCGCAACGCAATTAATGTGAGTTAGCTCACTCATTAGGCACCCCAGGCTTTACACTTTATGCTTCCGGCTCGTATGTTGTGTGGAATTGTGAGCGGATAACAATTTCACACAGGAAACAGCTATGACATGATTACGAATTAATTCGAGCTCGCCC
