## Supplementary material for "Pivotal role of Disrupted-in-Schizophrenia 1 (DISC1) in cardiac resilience to ischemic stress": Merged supplementary Figures S1-S6 Data S1. Tables S1-S4

### This PDF file includes:

Figures S1-S6
Data S1.
Tables S1-S4
References (1 to 2)

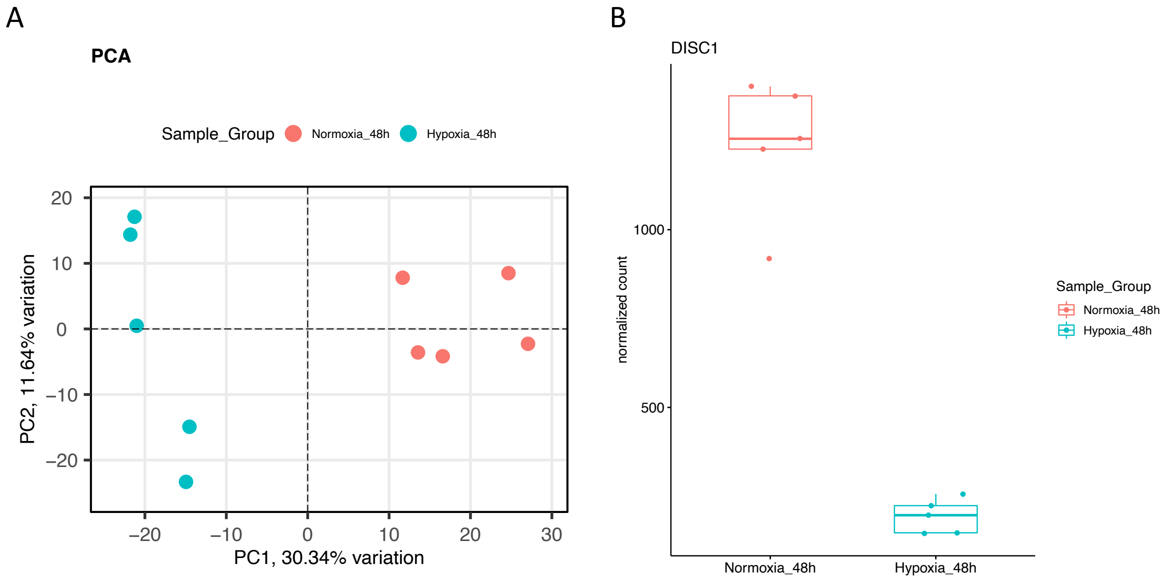

**Figure S1: RNA seq data:**

A) First two components of a Principal Component Analysis, with percentages of variance associated with each axis. B) expression of DISC1 in RNA seq data presented with box-whiskers including individual replicates.

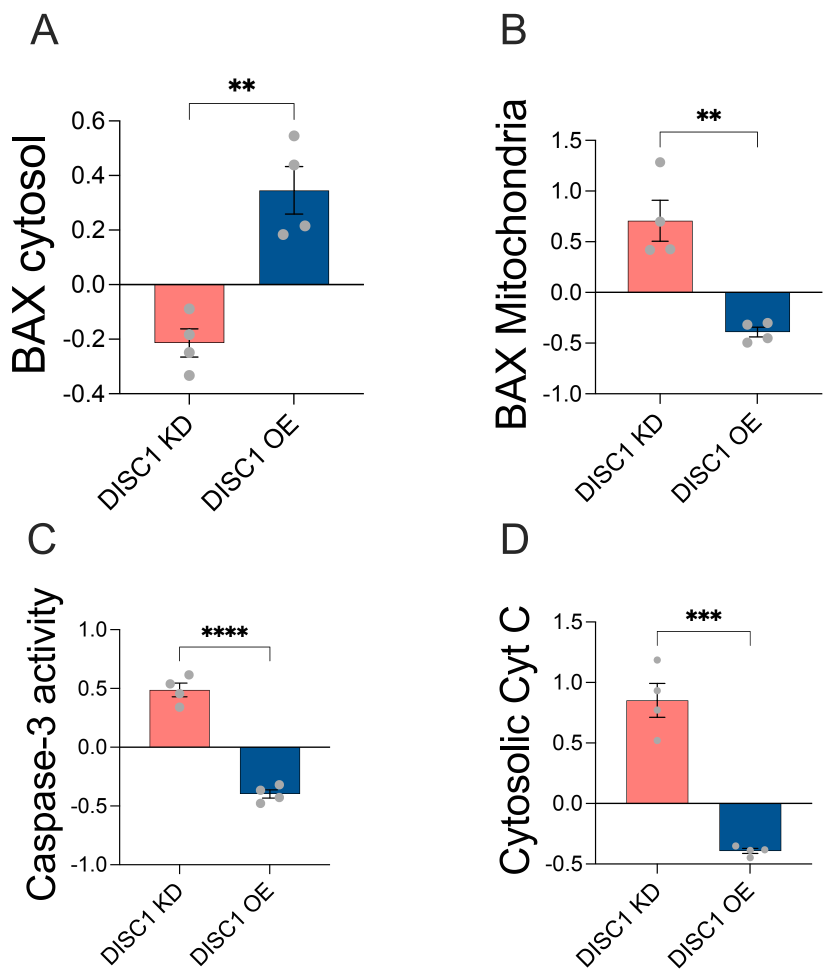

**Figure S2. Reduced DISC1 protein levels translocate BAX from cytosol to mitochondria.** Figures **A-B** represent experiments in in AC16 cardiomyocytes to determine the effect of both DISC1 silencing using short hairpin (sh)RNA pLKO.1-DISC1 (**DISC1 KD**) compared to respective pLKO.1- empty vector or by overexpression of DISC1 using pRK5-DISC1 overexpression vector (**DISC1 OE**) compared to respective pRK5-EV. All data are presented in relation to the fractional difference from respective Mock (zero line). **A**, BAX levels measured in isolated cytosolic fraction; **B**, BAX levels in isolated mitochondrial fractions; **C**, Cytosolic cytochrome C (Cyt C); **D**, caspase-3 activity. ELISA immunoassays validated the integrity and successful isolation of the distinguished components of the mitochondrial fraction and cytosolic fraction using the control proteins β-Actin and COX4 (Fig.S3). All data are presented as mean ± s.e.m. Statistical analyses in D-F were performed using two-tailed t-test to assess the difference in response compared to respective Mock between DISC1 KD versus DISC OE. Statistical difference displayed by: *** P<0.01*

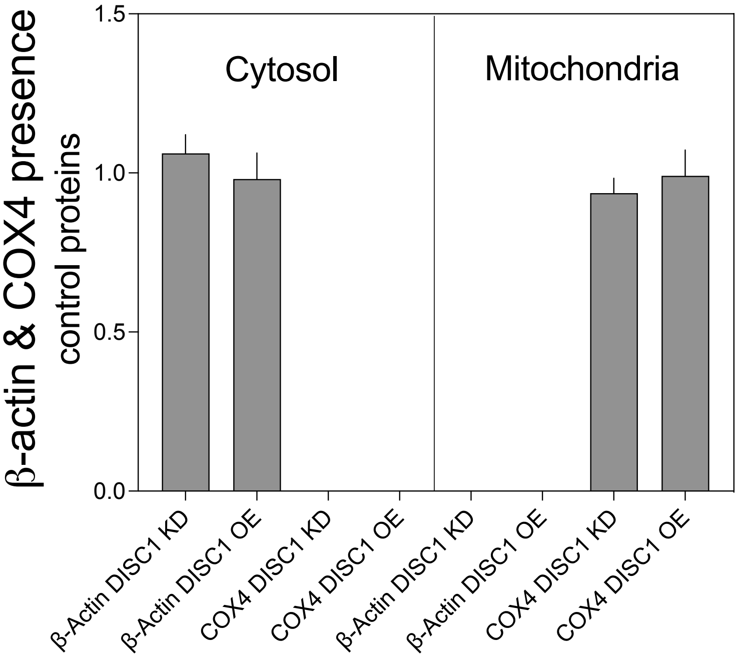

**Figure S3. Control experiments to verify cytosolic and mitochondrial fraction in data from figure S2 and figure S4.** ELISA immunoassays validated the integrity and successful isolation of the distinguished components of the mitochondrial fraction and cytosolic fraction. Prescence of COX4 and no detection of β-Actin verified successful of mitochondria. Presence of β-Actin and no detection of COX4 verified cytosolic fraction.

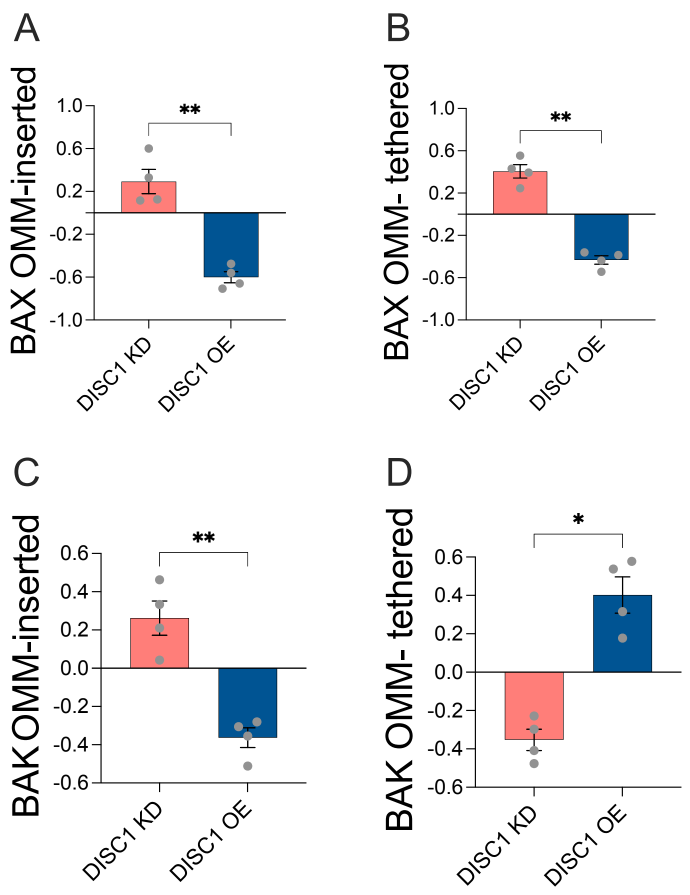

**Figure S4. Reduced DISC1 protein levels increase cell death during hypoxia by recruiting BAX from cytosol to integrate in the OMM and form MAC pores.** Figures **A-D** represent experiments in in AC16 cardiomyocytes to determine the effect of both DISC1 silencing using short hairpin (sh)RNA pLKO.1-DISC1 (**DISC1 KD**) compared to respective pLKO.1- empty vector or by overexpression of DISC1 using pRK5-DISC1 overexpression vector (DISC1 OE) compared to respective pRK5-EV. All data are presented in relation to the fractional difference from respective Mock (zero line). **A**, BAX in the outer mitochondria membrane (OMM)- inserted fraction; **B**, BAX in the OMM-tethered fraction; **C**, BAK in the OMM- inserted fraction; **D**, BAK in the OMM-tethered fraction. Statistical analyses in D-F were performed using two-tailed t-test to assess the difference in response compared to respective Mock between DISC1 KD versus DISC1 OE. Statistical difference displayed by: ** P<0.05; **P<0.01,* compared to respective Mock.

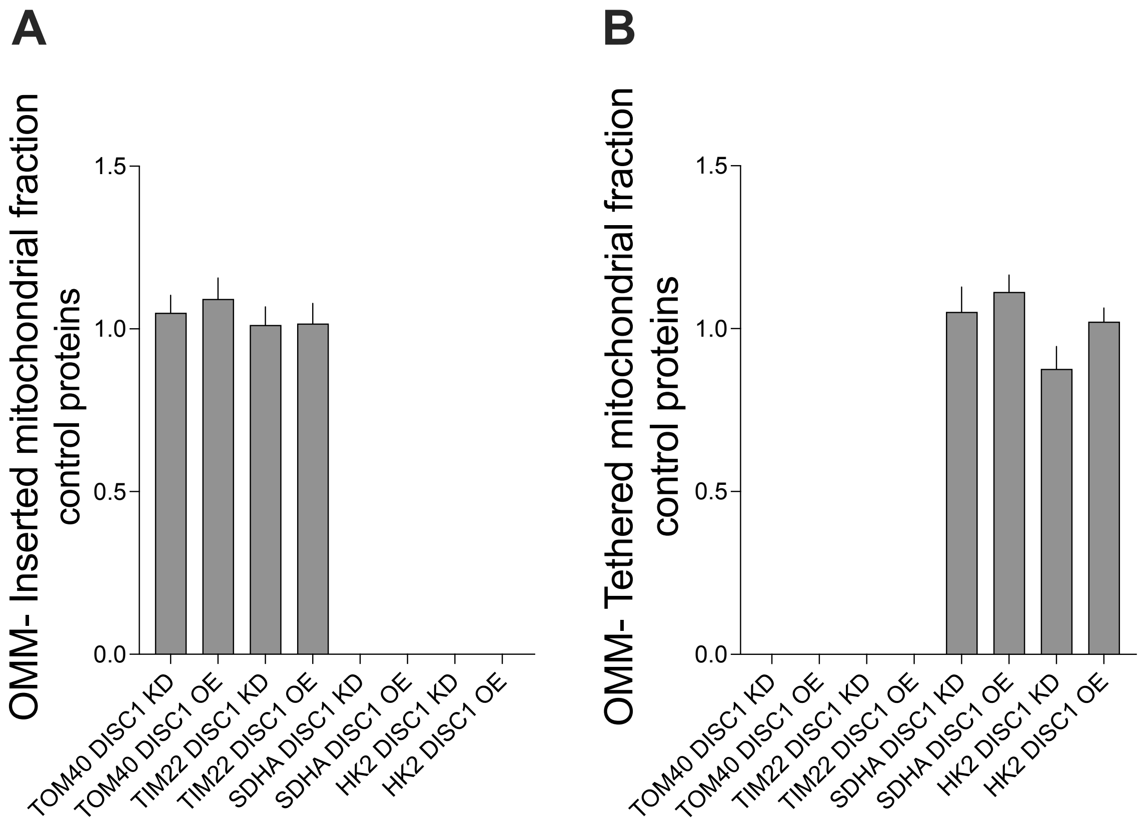

**Figure S5. Control experiments to verify and distinguish between the inserted and the tethered fraction of the** outer mitochondria membrane (OMM)**.** ELISA immunoassays validated the integrity and successful isolation of the distinguished components of the OMM. **A**, The OMM-inserted fraction was characterized by the presence of TOM40 (Mitochondrial import receptor subunit Translocase of outer membrane 40 kDa subunit) and TIM22(Translocase of inner mitochondrial membrane 22), but with absence of SDHA (Succinate Dehydrogenase) and HK2 (Hexokinase 2). **B**, The OMM-tethered fraction OMM-tethered fraction were absent of TOM40 and TIM22 concomitant with the presence of SDHA and HK2.

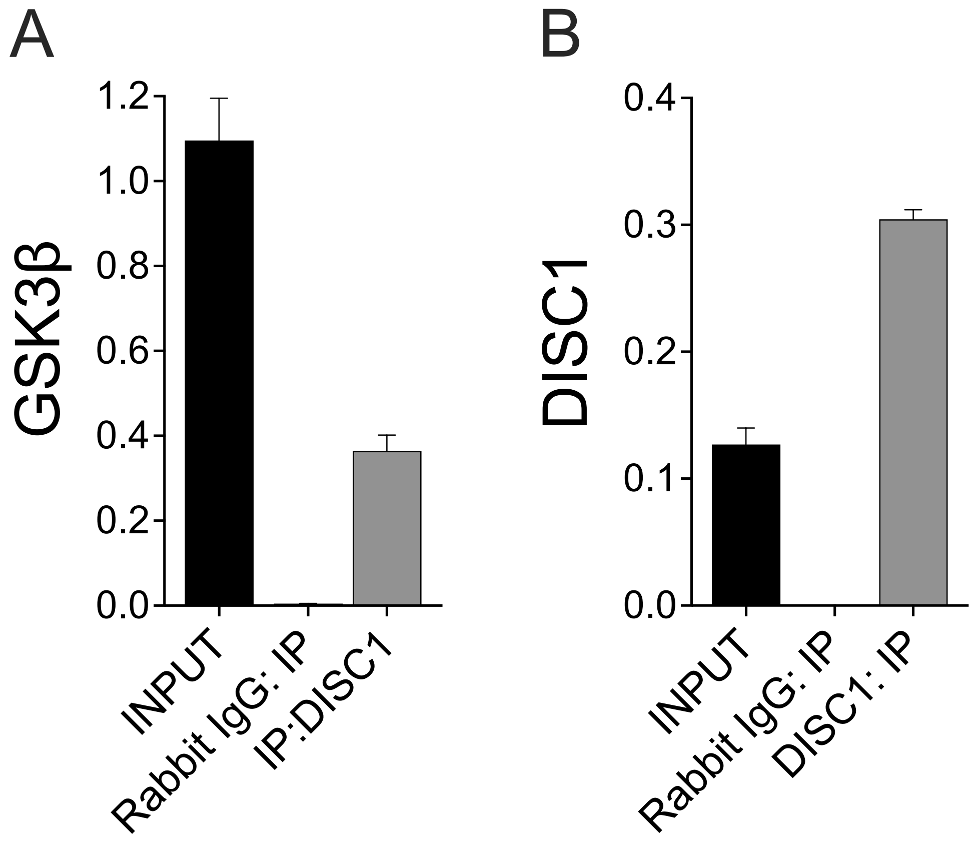

**Figure S6. Co-immunoprecipitation (Co-IP) coupled with tandem ELISA immunoassay determining the abundance of GSK3ß in the DISC1 immunoprecipitates, as a surrogate measure of the direct protein-protein interaction between DISC1 and GSK3ß.** (A) ELISA immunoassay showing the relative *quantitative abundance* of GSK3ß in the DISC1 immunoprecipitates compared to the control Rabbit IgG immunoprecipitates and the native *Input* (20%) fraction subjected to the DISC1 immunoprecipitation. (B) Quantitative ELISA immunoassay validating DISC1 immunoprecipitation and showing the abundance of DISC1 in the respective fractions. Data is expressed as experimental blank-corrected absorbances measured at $\lambda_{450}$ (450 nm). Data is expressed as mean ± S.D from three technical replicates for each of the four biological replicates.

**Data S1: GWAS enrichment analysis**

We performed genome-wide association study (GWAS) enrichment analysis to investigate whether risk genes for traits directly associated with cardiologic diseases, malformations, or pathologies (**Table S1**) are over-represented in DISC1 protein interactors [1]. Risk genes with 1 or more single nucleotide polymorphisms (SNP) associated with traits listed in **Table S1** (p-value < 1E-8 and odd ratio > 1) from the NHGRI-EBI GWAS Catalog[2] were used in the analysis. Over-representation was assessed using a chi-square test for the following 2×2 contingency table:

| | Risk Gene | Not Risk Gene |
|---|---|---|
| DISC1 protein interactors | 10 | 184 |
| Other genes which are not DISC1 protein interactors in human genome (GRCh38.84) | 298 | 58338 |

**Table S1. Traits directly associated with cardiologic diseases, malformations, or pathologies used in the GWAS enrichment analysis**

| Trait |
| --- |
| AL amyloidosis |
| Abdominal Aortic Aneurysm |
| Abnormal thrombosis |
| Aortic Coarctation |
| Brugada syndrome |
| D dimer measurement |
| Ischemic stroke |
| JT interval |
| P wave duration |
| P wave terminal force measurement |
| PR interval |
| PR segment |
| QRS amplitude |
| QRS complex |
| QRS duration |
| QT interval |
| RR interval |
| TPE interval measurement |
| acute coronary syndrome |
| aortic stenosis |
| aortic valve calcification |
| atrial fibrillation |
| brain aneurysm |
| cardiac embolism |
| cardiovascular event measurement |
| cervical artery dissection |
| chronic obstructive pulmonary disease |
| clinical ideal cardiovascular health |
| congenital heart disease |
| congenital heart malformation |
| congenital left-sided heart lesions |
| coronary artery disease |
| coronary heart disease |
| deep vein thrombosis |
| diarrhea |
| diastolic blood pressure |
| dilated cardiomyopathy |
| electrocardiography |
| heart amyloid deposition measurement |
| heart failure |
| heart function measurement |
| heart rate |
| idiopathic dilated cardiomyopathy |
| large artery stroke |
| mean arterial pressure |
| mortality |
| myocardial infarction |
| myositis |

peripheral arterial disease
pulmonary artery enlargement
pulmonary embolism
pulmonary hypertension
pulse pressure measurement
response to beta blocker
response to cold pressor test
response to darapladib
response to dietary potassium supplementation
response to high sodium diet
response to low sodium diet
response to sulfonylurea
response to tetracyclic antidepressant
response to tricyclic antidepressant
resting heart rate
rheumatic heart disease
small artery occlusion
small vessel stroke
stroke
sudden cardiac arrest
systolic blood pressure
temporal arteritis
tetralogy of fallot
thoracic aortic aneurysm
traffic air pollution measurement
vascular dementia
vasoactive peptide measurement
venous thromboembolism

| Table S2. Patients characteristics | |
|---|---|
| ***Baseline characteristics*** | |
| Gender, men/women, n | 22/7 |
| Age, years | 68 ± 8 |
| Height, cm | 173,1 ± 8,8 |
| Weight, kg | 83,1 ± 15 |
| Body mass index, kg/m$^2$ | 27,7 ± 4 |
| ASA score | 3.6 ± 0.6 |
| ***Risk factors and comorbidities, n*** | |
| Chronic obstructive pulmonary disease | 2 |
| Current smoker | 8 |
| History of smoking | 21 |
| Diabetes mellitus | 5 |
| Hypertension | 14 |
| Peripheral arterial disease | 1 |
| Previous cerebral insult | 0 |
| Previous myocardial infarction | 15 |
| Previous PCI | 5 |
| ***Pharmacotherapy, n*** | |
| ACE-inhibitor or ARB | 10 |
| Aspirin | 28 |
| Beta blocker | 24 |
| Calcium channel blocker | 8 |
| Clopidogrel | 11 |
| Dipyridamole | 0 |
| Diuretics | 4 |
| Glibenclamide | 1 |
| Insulin | 1 |
| Lipid-lowering agent | 26 |
| Metformin | 3 |
| Organic nitrates | 6 |
| Sulfonylurea | 0 |
| Warfarin | 2 |
| ***Intraoperative data*** | |
| Aortic cross-clamping, min | 42 ± 11 |
| Cardiopulmonary bypass, min | 77 ± 23 |
| Cardioplegia, mL | 1003 ± 288 |
| Distal coronary graft anastomoses, n | 3,2 ± 0.7 |

**Table S2 Notation legend:** Clinical data from CABG patients supporting figure 3D.
Data are presented as mean ± SD or number. ASA-score refers to the preoperative physical assessment score developed by the American Society of Anesthesiology. PCI stands for percutaneous coronary intervention. ACE denotes angiotensin converting enzyme, and ARB represents angiotensin-II-receptor blocker.

### Table S3 - Composition of the *hypoxia medium*

| Component | 500 mL | Final concentration | Source (Notation) |
|---|---|---|---|
| DMEM, No Glucose | 464.45 mL | 93% v/v | 1 |
| Creatine | 131.2 mg | 5 mM | 2 |
| D-(+)-Glucose Solution 2.5 M, 450 g/L | 0.55 mL | 2.75 mM | 3 |
| Glutamine 200 Mm | 5 mL | 2 mM | 4 |
| HEPES 1M | 5 mL | 10 mM | 5 |
| L-Carnitine, 200 mM | 5 mL | 2 mM | 6 |
| Non-essential Amino Acids, 100x | 5 mL | N/A* | 7 |
| Sodium Pyruvate 100 mM | 5 mL | 1 mM | 8 |
| Taurine 500 mM | 5 mL | 5 mM | 9 |
| Linoleic Acid-Oleic Acid-Albumin, 100x | 5 mL | N/A* | 10 |

**Table S3 Notation legend:**

[1] Thermo Fisher Scientific, Oslo, Norway, Catalogue # 11966025
[2] Sigma Aldrich / Merck Millipore / Merck Life Science, Darmstadt, Germany, Catalogue # C3630-100G
[3] Sigma Aldrich / Merck Millipore / Merck Life Science, Oslo, Norway, Catalogue # G8769
[4] Thermo Fisher Scientific, Oslo, Norway, Catalogue # A2916801
[5] Sigma Aldrich / Merck Millipore / Merck Life Science, Darmstadt, Germany, Catalogue # H4034-500G
[6] Sigma Aldrich / Merck Millipore / Merck Life Science, Darmstadt, Germany, Catalogue # C0283-25G
[7] Thermo Fisher Scientific, Oslo, Norway, Catalogue # 11140035
[8] Thermo Fisher Scientific, Oslo, Norway, Catalogue # 11360070
[9] Sigma Aldrich / Merck Millipore / Merck Life Science, Darmstadt, Germany, Catalogue # T8691-100G
[10] Sigma Aldrich / Merck Millipore / Merck Life Science, Darmstadt, Germany, Catalogue # L9655-5ML
* N/A - Not Applicable

### Table S4 - List of monoclonal and polyclonal antibodies used in the study

| Antibody | Application | Amount | Host | Manufacturer | Catalogue # | Resource Identifier ID (RRID) |
|---|---|---|---|---|---|---|
| β-Actin | ELISA *Capture* | 20 ng / well | Mouse | Santa Cruz Biotechnology | sc-47778 | AB_2714189 |
| β-Actin | ELISA *Detection* | 20 ng / well | Rabbit | Cell Signaling Technology | 4970 | AB_2223172 |
| β-Actin antibody blocking peptide | ELISA *Detection* | N/A | N/A | Cell Signaling Technology | 1025 | N/A |
| BAK | ELISA *Capture* | 20 ng / well | Mouse | Thermo Fisher Scientific | MA5-36225 | AB_2884059: |
| BAK | ELISA *Detection* | 20 ng / well | Rabbit | Novus Biologicals | NBP1-77152 | AB_11014847 |
| BAK antibody blocking peptide | ELISA *Detection* | N/A | N/A | Novus Biologicals | NBP1-77152PEP | N/A |
| BAX | ELISA *Capture* | 20 ng / well | Mouse | Thermo Fisher Scientific | 33-6600 | AB_2533133 |
| BAX | ELISA *Detection* | 20 ng / well | Rabbit | Novus Biologicals | NBP1-88682 | AB_11014342 |
| BAX antibody blocking peptide | ELISA *Detection* | N/A | N/A | Novus Biologicals | NBP1-88682PEP | N/A |
| COX4 | ELISA *Capture* | 20 ng / well | Mouse | Thermo Fisher Scientific | MA5-15686 | AB_10977841 |
| COX4 | ELISA *Detection* | 20 ng / well | Rabbit | Cell Signaling Technology | 4844 | AB_2085427 |
| COX4 antibody blocking peptide | ELISA *Detection* | N/A | N/A | Cell Signaling Technology | 1034 | N/A |
| Cytochrome C | ELISA *Capture* | 20 ng / well | Mouse | Thermo Fisher Scientific | BMS1037 | AB_10598651 |
| Cytochrome C | ELISA *Detection* | 20 ng / well | Rabbit | Cell Signaling Technology | 4280 | AB_10695410 |
| DISC1 | ELISA *Capture* | 10 ng / well | Mouse | R&D Systems | MAB6699 | AB_10890920 |
| DISC1 | ELISA *detection* | 10 ng / well | Rabbit | Thermo Fisher Scientific | PA5-20422 | AB_11153106 |
| Goat Anti-Mouse IgG (H + L)-HRP Conjugate | 1:5000 | 1 µg | Goat | Bio-Rad | 1706516 | AB_11125547 |
| Goat Anti-Mouse IgG-AP Conjugate | 1:5000 | N/A € | Goat | Bio-Rad | 1706520 | AB_11125348 |
| Goat Anti-Rabbit IgG (H + L)-HRP Conjugate | 1:5000 | 1 µg | Goat | Bio-Rad | 1706515 | AB_11125142 |
| Goat Anti-Rabbit IgG-AP Conjugate | 1:20000 | N/A € | Goat | Sigma Aldrich / Merck Life Science | A3687 | AB_258103 |
| GSK3β | IP | 5 µg / IP | Rabbit | Cell Signaling Technology | 9315 | AB_490890 |
| GSK3β | ELISA *Capture* | 10 ng / well | Mouse | Cell Signaling Technology | 9832 | AB_10839406 |
| GSK3β | ELISA *Detection* | 10 ng / well | Rabbit | Cell Signaling Technology | 9315 | AB_490890 |
| GSK3β antibody blocking peptide | ELISA *Detection* | N/A | N/A | Cell Signaling Technology | 1073 | N/A |
| HK2 | ELISA *Capture* | 30 ng / well | Rabbit | Thermo Fisher Scientific | PA5-97828 | AB_2812442 |
| HK2 | ELISA *Detection* | 30 ng / well | Mouse | Thermo Fisher Scientific | MA5-15679 | AB_10986812 |
| LDH | ELISA *Capture* | 30 ng / well | Mouse | Santa Cruz Biotechnology | sc-133123 | AB_2134964 |
| LDH-A | ELISA *Detection* | 30 ng / well | Rabbit | Novus Biologicals | NBP1-48336 | AB_10011099 |
| LDH-A antibody blocking peptide | ELISA *Detection* | N/A | N/A | Novus Biologicals | NBP1-48336PEP | N/A |
| LDH-B | ELISA *Detection* | 30 ng / well | Rabbit | Novus Biologicals | NBP2-38131 | N/A |

| | | | | | | |
|---|---|---|---|---|---|---|
| LDH-B antibody blocking peptide | ELISA *Detection* | N/A | N/A | Novus Biologicals | NBP2-38131PEP | N/A |
| p-Ser | GSK3β activity | 100 ng /well | Mouse | Santa Cruz Biotechnology | sc-81516 | AB_1128626 |
| Rabbit IgG | IP | 5 µg / IP | Rabbit | Santa Cruz Biotechnology | sc-3888 | AB_737196 |
| SDHA (SDH2) | ELISA *Capture* | 30 ng / well | Mouse | Thermo Fisher Scientific | 459200 | AB_2532231 |
| SDHA (SDH2) | ELISA *detection* | 30 ng / well | Rabbit | Cell Signaling Technology | 11998 | AB_2750900 |
| TOM40 | ELISA *capture* | 30 ng / well | Mouse | Santa Cruz Biotechnology | sc-365467 | AB_10847086 |
| TOM40 | ELISA *detection* | 30 ng / well | Rabbit | Thermo Fisher Scientific | 18409-1-AP | AB_2303725 |
| TIM22 | ELISA *capture* | 30 ng / well | Mouse | Sigma Aldrich / Merck Life Science | SAB1400520- | AB_1858016 |
| TIM22 | ELISA *detection* | 30 ng / well | Rabbit | Thermo Fisher Scientific | 14927-1-AP | AB_11183050 |

**N/A:** Not Available / Not Applicable
IP: Immunoprecipitation
**€:** Amount of secondary antibody cannot be determined as the commercial vendor does not provide the antibody concentration
